# Unlocking DNA Damage Sensitivity of Cancer Cells: The Potential of Splicing Inhibitors

**DOI:** 10.1101/2023.10.08.561421

**Authors:** Ksenia S. Anufrieva, Maria M. Lukina, Olga M. Ivanova, Anastasia N. Kazakova, Polina V. Shnaider, Ksenia M. Klimina, Vladimir A. Veselovsky, Artem V. Luzhin, Artem K. Velichko, Omar L. Kantidze, Elizaveta N. Mochalova, Maxim P. Nikitin, Aleksandra V. Kashina, Ekaterina A. Vasilchikova, Roman V. Deev, Alexey M. Emelin, Anton N. Turchin, Zhaojian Liu, Zixiang Wang, Veronika S. Boichenko, Nadezhda M. Markina, Maria A. Lagarkova, Vadim M. Govorun, Georgij P. Arapidi, Victoria O. Shender

## Abstract

Despite the growing interest in pre-mRNA alternative splicing (AS) as a therapeutic anticancer target, the potential of splicing inhibitors in treating solid tumors remains largely unexplored. We conducted a meta-analysis of transcriptome data from six different tumor types and revealed that splicing inhibitors induced similar patterns of AS, resulting in widespread exon-skipping and intron retention events that often lead to nonsense-mediated decay of the transcripts. Interestingly, in many cases exon skipping is induced by a compensatory cellular response to splicing inhibitor treatment. It involves an upregulation of multiple splicing factors and incomplete recognition of branch points by U2 snRNP. These post transcriptional changes downregulate one-third of essential DNA repair genes, thereby creating a therapeutic vulnerability that can be exploited for cancer treatment. To harness this vulnerability, we proposed a new approach to cancer treatment consisting of sequential addition of a splicing inhibitors followed by a DNA-damaging agent. Our *in vitro* and *in vivo* experiments demonstrated that this strategy exhibits promising therapeutic potential for a wide range of tumors.

## INTRODUCTION

Transcriptome analyses have highlighted that aberrant pre-mRNA splicing is a characteristic feature across various cancer types, even in the absence of splicing factor mutations^1^. As a result, the spliceosomal machinery is emerging as a promising target for innovative therapeutic interventions^2–4^. Diverse strategies, encompassing small molecule inhibitors to oligonucleotide-based approaches, have been proposed^5,6^. While oligonucleotide-based therapy offers heightened specificity, the substantial heterogeneity in malignant tumors poses a challenge. Small molecule-based splicing inhibitors, including drugs targeting spliceosome assembly at various stages (early: spliceostatins, pladienolides, herboxidienes; late: isogingketin), along with those impeding post-translational modifications of spliceosomal proteins, antibiotics, and sulfonamides, present viable options. Notably, inhibitors targeting the early stage of spliceosome assembly demonstrate significant anticancer cytotoxicity both *in vitro* and *in vivo*, and pladienolides and their synthetic analogues (E7107 and H3B-8800), are recognized as the most potent splicing inhibitors^6^.

Clinical trials of E7107 against several solid tumors were halted due to dose-limiting toxicity for patients^7,8^, while the therapeutic potential of H3B-8800 is being assessed in phase I/II trials (NCT02841540) against myeloid neoplasms with splicing factor mutations, considered to confer synthetic lethality to spliceosome inhibitors^9,10^. While both compounds disrupt pre-mRNA splicing and inhibit tumor growth, complete tumor elimination remains elusive. Despite the current limitations of targeting the spliceosome in monotherapy, exploring drug combinations may pave the way for innovative splicing-based cancer treatments. A comprehensive understanding of splicing inhibitors’ mechanism of action is crucial for devising effective therapeutic strategies.

It is assumed that the action of these pladienolide-derived inhibitors is centered on binding to the spliceosomal protein SF3B1, a key component of the U2 small nuclear ribonucleoprotein complex (U2 snRNP), pivotal in the early stages of splicing site recognition^11^. This leads to less stable interactions of U2 snRNP with pre-mRNA sequences, resulting in substantial changes in pre-mRNA alternative splicing^12–14^. However, consensus on the mechanisms mediating cancer cell death remains elusive. Splicing inhibitors have been linked to the production of pro-apoptotic mRNA isoforms of genes such as *BAX*, *BCL2L1^13,15^*. Other studies propose that the blockade of SF3B1 activity induces excessive toxic R-loops^16,17^, while further investigations suggest a reduction in RNA synthesis, caused by prolonged RNA Polymerase II promoter-proximal pausing and a slower elongation rate at the beginning of genes^17,18^. On the other hand, it was shown that splicing inhibitors cause exon-skipping changes which affect transcripts of DNA repair genes, such as *CHEK2^16,19^*. The latter mechanism is in good agreement with the fact that spliceosomal proteins play an important role in preventing DNA damage through their splicing-dependent or independent functions^20–24^. Additionally, the extensive study of splicing inhibitors on leukemia cell lines poses challenges in comprehensively understanding general molecular mechanisms partly due to the high frequency of splicing factor mutations in this malignancy.

Therefore, this study conducts a meta-analysis of transcriptome data from eight cancer cell lines representing six tumor types to identify common changes in gene expression and pre-mRNA splicing induced by splicing inhibitors. Regardless of tumor type, splicing inhibitors were observed to reduce the expression of approximately 94 DNA repair genes through exon skipping-triggered nonsense-mediated decay (NMD). Our proteomic and transcriptomic time series analysis revealed that NMD-associated exon skippings are a compensatory cellular response, accompanied by a significant increase in the expression of numerous splicing factors. Capitalizing on the DNA repair vulnerability caused by splicing inhibitors, we propose a novel cancer treatment approach involving the sequential addition of a splicing inhibitor and a DNA-damaging agent, opening up extensive possibilities for their future practical applications in tumor treatment.

## RESULTS

### Treatment with splicing inhibitors induces common pre-mRNA splicing and gene expression alterations across diverse cancer cell lines

In order to unravel the molecular mechanisms underpinning the potent antitumor activity of splicing inhibitors, we conducted an analysis utilizing publicly available RNA sequencing (RNA-seq) datasets from various cancer cell lines, including prostate adenocarcinoma (LNCaP, PC3), colorectal cancer (HCT-116), multiple myeloma (MM.1S, AMO-1), breast adenocarcinoma (MDA-MB-468), and T-cell leukemia (CUTLL1), which were treated with the same splicing inhibitor E7107 (Fig. 1A, Table 1). Initially, we scrutinized the transcriptomic changes induced by E7107 in each cancer cell line. E7107 treatment prompted approximately 29,000 alternative splicing events in every cell line, with 60-70% of these events being exon skippings (Fig. 1B, Extended Data Fig. 1A, Supplementary Table S1). Intriguingly, exposure to E7107 not only resulted in substantial changes in pre-mRNA splicing but also led to a notable decrease in gene expression (Fig. 1C). Functional analysis of alternatively spliced and downregulated genes revealed that E7107 disrupts the regulation of such pathways as cell cycle, DNA repair, and DNA replication (Fig. 1D, Extended Data Fig. 1B). Furthermore, our analysis demonstrated a significant intersection between the downregulation of gene expression and the exon skipping events induced by E7107 in each cell line (Fig. 1E). These findings suggest that the aberrant pre-mRNA splicing induced by the E7107 inhibitor makes transcripts more susceptible to degradation.

**Figure 1.**
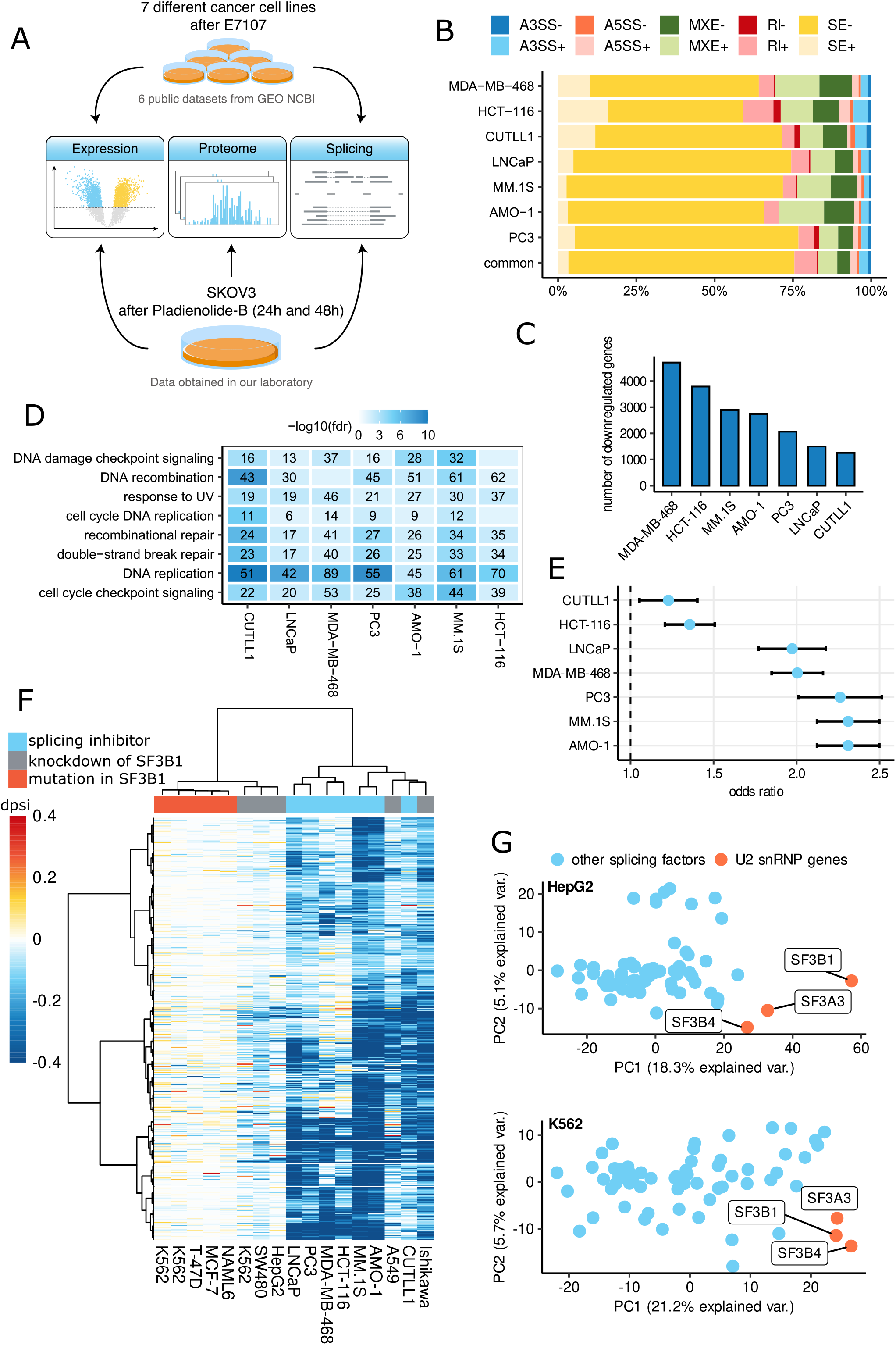
Treatment with splicing inhibitors induces common pre-mRNA splicing and gene expression alterations across diverse cancer cell lines. **A** – Overview of the analysis of RNA-seq and LS-MS/MS data from cancer cell lines treated with splicing inhibitors. **B** – Stacked bar plot illustrating the percentage of each type of alternative splicing event among all differential splicing events in 7 different cancer cell lines after E7107 treatment. Splicing events: SE - skipped exon, A5SS - alternative 5′ splice site, A3SS - alternative 3′ splice site, RI - retained intron, MXE - mutually exclusive exons. The plus sign after the event abbreviation denotes differential splicing events with dpsi > 0.05, and the minus sign denotes events with dpsi < −0.05. **C** – Bar plot indicating the number of downregulated genes for each cancer cell line after E7107 treatment. Changes in expression were considered significant if fdr<0.05 and log2-fold change < −1. **D** – Heat map displaying Reactome pathway enrichment analysis, showcasing common pathways downregulated in 7 cancer cell lines after E7107 treatment. The x-axis denotes the names of cancer cell lines, while the y-axis shows the pathway names. The numbers in the boxes represent the count of genes associated with the specific pathway for the given cancer cell line, with FDR < 0.05. The blue color scale reflects the false discovery rate (FDR). **E** – Graph presenting odds ratios and 95% confidence intervals, comparing the overlaps between skipped exons and differentially downregulated genes in 7 different cancer cell lines after E7107 treatment. All reported associations were significant at a level below 0.00001, with p-values calculated using Fisher’s exact test. **F** – Heat map illustrating splicing inhibitor-induced exon skipping events. Rows represent tumor-independent exon skipping events induced by splicing inhibitor treatment, and columns represent cell lines after splicing inhibitor treatment/*SF3B1* knockdown/*SF3B1* mutation. Details about each original RNA-seq experiment can be found in Table 1. Heatmap matrix values indicate dpsi values for each RNA-seq experiment. **G** – Principal Component Analysis (PCA) based on dpsi values of exon skipping events detected after *SF3B1* knockdown in HepG and K562 cell lines. Each point represents the alternative splicing profile of cells with the knockdown of one of the splicing factors in HepG and K562 cell lines. All data were obtained through the analysis of publicly available datasets from the ENCODE project.

**Table 1.**
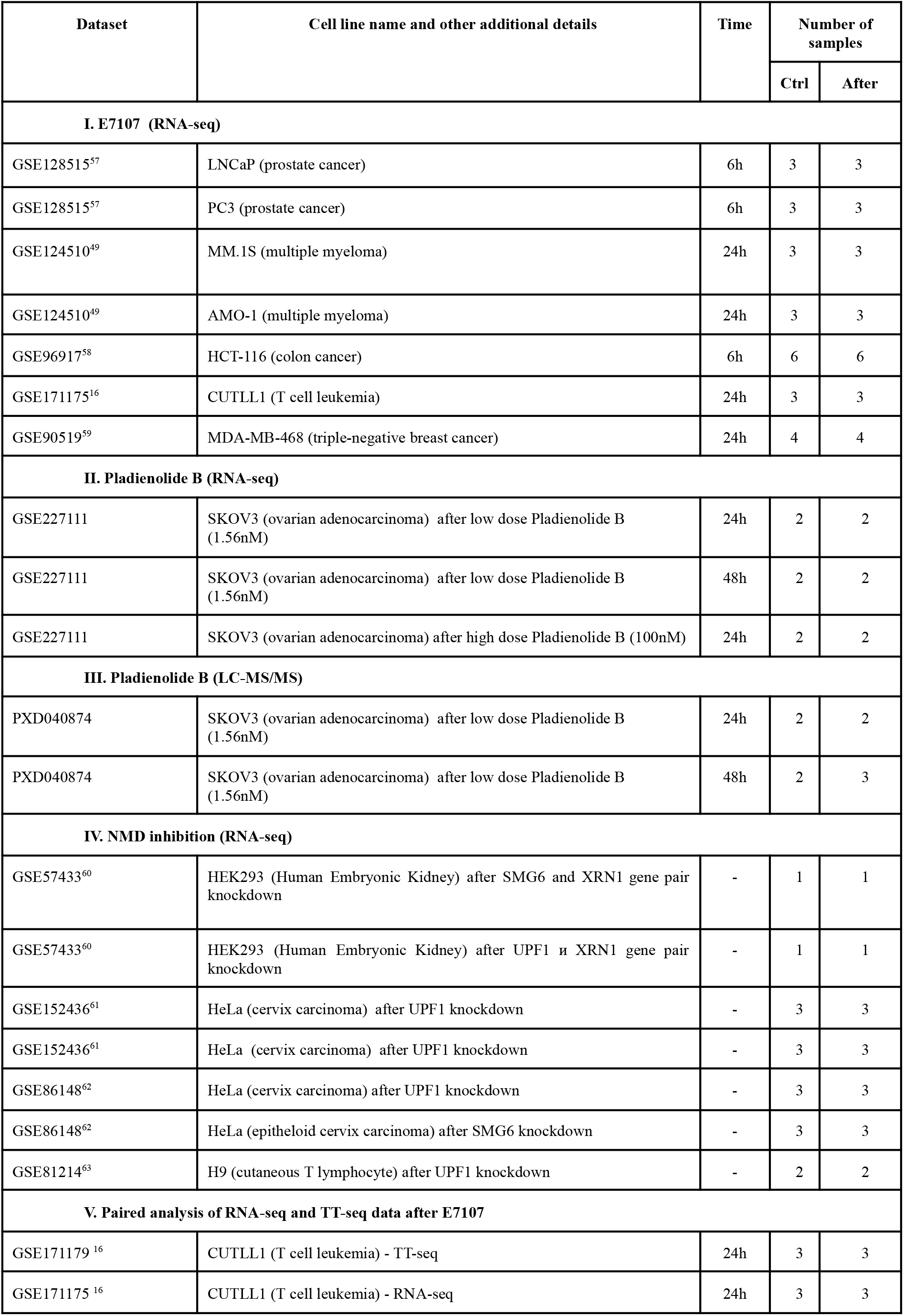
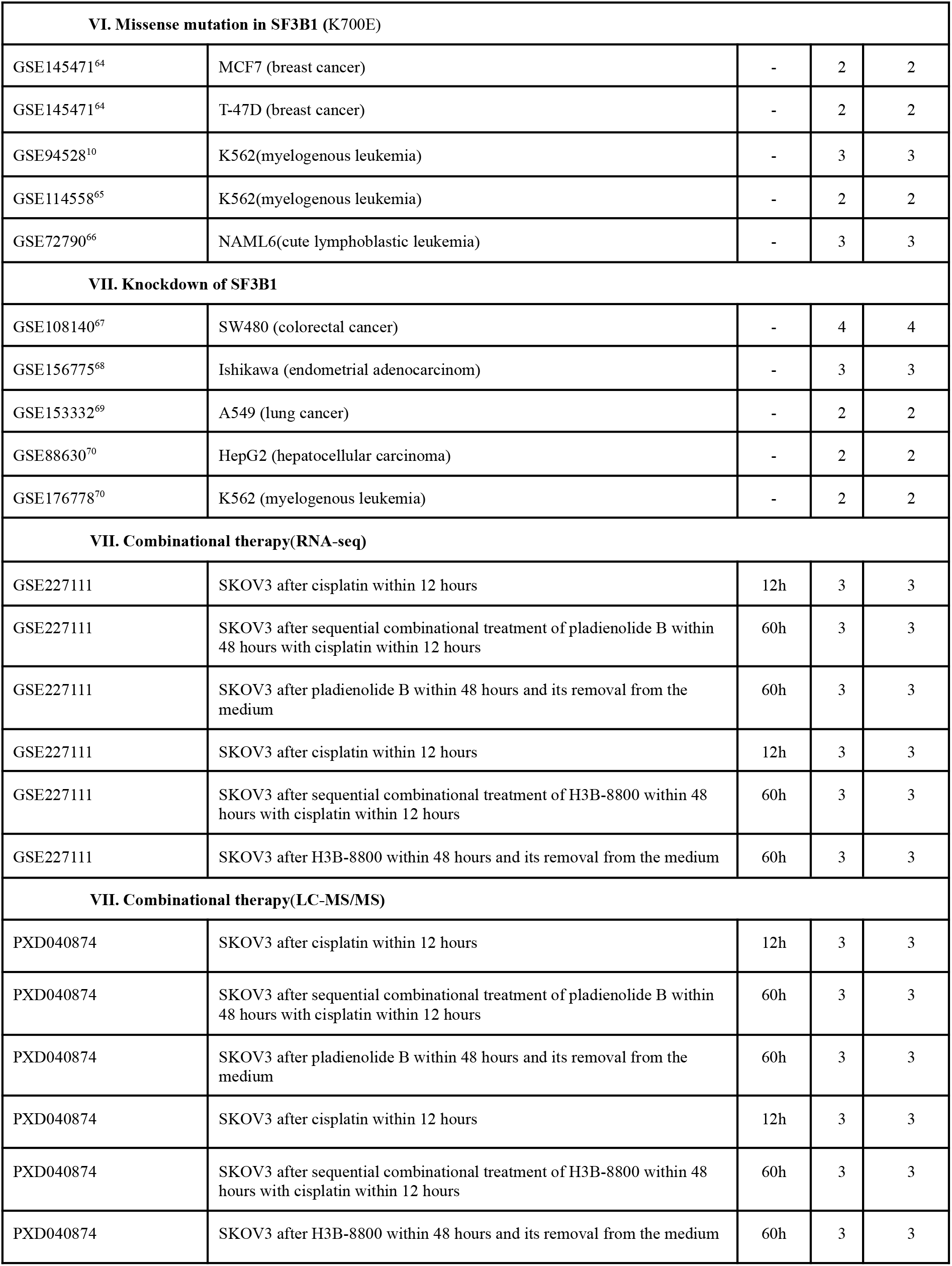
The RNA-Seq, TT-seq, LC-MS/MS data used in this study.

| Dataset | Cell line name and other additional details | Time | Number of samples |  |
| --- | --- | --- | --- | --- |
|  |  |  | Ctrl | After |
| I. E7107 (RNA-seq) |  |  |  |  |
| GSE128515<sup>57</sup> | LNCaP (prostate cancer) | 6h | 3 | 3 |
| GSE128515<sup>57</sup> | PC3 (prostate cancer) | 6h | 3 | 3 |
| GSE124510<sup>49</sup> | MM.1S (multiple myeloma) | 24h | 3 | 3 |
| GSE124510<sup>49</sup> | AMO-1 (multiple myeloma) | 24h | 3 | 3 |
| GSE96917<sup>58</sup> | HCT-116 (colon cancer) | 6h | 6 | 6 |
| GSE171175<sup>16</sup> | CUTLL1 (T cell leukemia) | 24h | 3 | 3 |
| GSE90519<sup>59</sup> | MDA-MB-468 (triple-negative breast cancer) | 24h | 4 | 4 |
| II. Pladienolide B (RNA-seq) |  |  |  |  |
| GSE227111 | SKOV3 (ovarian adenocarcinoma) after low dose Pladienolide B (1.56nM) | 24h | 2 | 2 |
| GSE227111 | SKOV3 (ovarian adenocarcinoma) after low dose Pladienolide B (1.56nM) | 48h | 2 | 2 |
| GSE227111 | SKOV3 (ovarian adenocarcinoma) after high dose Pladienolide B (100nM) | 24h | 2 | 2 |
| III. Pladienolide B (LC-MS/MS) |  |  |  |  |
| PXD040874 | SKOV3 (ovarian adenocarcinoma) after low dose Pladienolide B (1.56nM) | 24h | 2 | 2 |
| PXD040874 | SKOV3 (ovarian adenocarcinoma) after low dose Pladienolide B (1.56nM) | 48h | 2 | 3 |
| IV. NMD inhibition (RNA-seq) |  |  |  |  |
| GSE57433<sup>60</sup> | HEK293 (Human Embryonic Kidney) after SMG6 and XRN1 gene pair knockdown | - | 1 | 1 |
| GSE57433<sup>60</sup> | HEK293 (Human Embryonic Kidney) after UPF1 и XRN1 gene pair knockdown | - | 1 | 1 |
| GSE152436<sup>61</sup> | HeLa (cervix carcinoma) after UPF1 knockdown | - | 3 | 3 |
| GSE152436<sup>61</sup> | HeLa (cervix carcinoma) after UPF1 knockdown | - | 3 | 3 |
| GSE86148<sup>62</sup> | HeLa (cervix carcinoma) after UPF1 knockdown | - | 3 | 3 |
| GSE86148<sup>62</sup> | HeLa (epitheloid cervix carcinoma) after SMG6 knockdown | - | 3 | 3 |
| GSE81214<sup>63</sup> | H9 (cutaneous T lymphocyte) after UPF1 knockdown | - | 2 | 2 |
| V. Paired analysis of RNA-seq and TT-seq data after E7107 |  |  |  |  |
| GSE171179 <sup>16</sup> | CUTLL1 (T cell leukemia) - TT-seq | 24h | 3 | 3 |
| GSE171175 <sup>16</sup> | CUTLL1 (T cell leukemia) - RNA-seq | 24h | 3 | 3 |

| VI. Missense mutation in SF3B1 (K700E) |  |  |  |  |
| --- | --- | --- | --- | --- |
| GSE145471<sup>64</sup> | MCF7 (breast cancer) | - | 2 | 2 |
| GSE145471<sup>64</sup> | T-47D (breast cancer) | - | 2 | 2 |
| GSE94528<sup>10</sup> | K562(myelogenous leukemia) | - | 3 | 3 |
| GSE114558<sup>65</sup> | K562(myelogenous leukemia) | - | 2 | 2 |
| GSE72790<sup>66</sup> | NAML6(cute lymphoblastic leukemia) | - | 3 | 3 |
| VII. Knockdown of SF3B1 |  |  |  |  |
| GSE108140<sup>67</sup> | SW480 (colorectal cancer) | - | 4 | 4 |
| GSE156775<sup>68</sup> | Ishikawa (endometrial adenocarcinoma) | - | 3 | 3 |
| GSE153332<sup>69</sup> | A549 (lung cancer) | - | 2 | 2 |
| GSE88630<sup>70</sup> | HepG2 (hepatocellular carcinoma) | - | 2 | 2 |
| GSE176778<sup>70</sup> | K562 (myelogenous leukemia) | - | 2 | 2 |
| VII. Combinational therapy(RNA-seq) |  |  |  |  |
| GSE227111 | SKOV3 after cisplatin within 12 hours | 12h | 3 | 3 |
| GSE227111 | SKOV3 after sequential combinational treatment of pladienolide B within 48 hours with cisplatin within 12 hours | 60h | 3 | 3 |
| GSE227111 | SKOV3 after pladienolide B within 48 hours and its removal from the medium | 60h | 3 | 3 |
| GSE227111 | SKOV3 after cisplatin within 12 hours | 12h | 3 | 3 |
| GSE227111 | SKOV3 after sequential combinational treatment of H3B-8800 within 48 hours with cisplatin within 12 hours | 60h | 3 | 3 |
| GSE227111 | SKOV3 after H3B-8800 within 48 hours and its removal from the medium | 60h | 3 | 3 |
| VII. Combinational therapy(LC-MS/MS) |  |  |  |  |
| PXD040874 | SKOV3 after cisplatin within 12 hours | 12h | 3 | 3 |
| PXD040874 | SKOV3 after sequential combinational treatment of pladienolide B within 48 hours with cisplatin within 12 hours | 60h | 3 | 3 |
| PXD040874 | SKOV3 after pladienolide B within 48 hours and its removal from the medium | 60h | 3 | 3 |
| PXD040874 | SKOV3 after cisplatin within 12 hours | 12h | 3 | 3 |
| PXD040874 | SKOV3 after sequential combinational treatment of H3B-8800 within 48 hours with cisplatin within 12 hours | 60h | 3 | 3 |
| PXD040874 | SKOV3 after H3B-8800 within 48 hours and its removal from the medium | 60h | 3 | 3 |

Following that, we explored the consistent patterns of gene expression and pre-mRNA splicing induced by E7107 treatment across various tumor types (Supplementary Table S1-2). Remarkably, in at least half of the cell lines, E7107 induced alterations in the splicing of 4,348 genes and downregulated the expression of 1,441 genes (Extended Data Fig. 1C). Enrichment analysis further demonstrated that the commonly downregulated and alternatively spliced genes were associated with the DNA repair pathway (Extended Data Fig. 1D-E).

Our investigation unveiled that the splicing inhibitor targeting SF3B1 primarily diminishes the expression of DNA repair genes. Nonetheless, it is recognized that SF3B1 can also undergo dysregulation through mutations, frequently found in patients with chronic lymphocytic leukemia and myelodysplastic syndromes^25,26^. Analyzing publicly available RNA-seq data from four cancer cell lines with and without the hotspot K700E *SF3B1* mutation, we observed that the mutation alone could not replicate the splicing changes observed under the influence of splicing inhibitors (Table 1, Fig. 1F, Supplementary Table S1). However, the analysis of transcriptomic profiles in five different cancer cell lines with SF3B1 knockdown exhibited similar patterns of gene expression and pre-mRNA splicing as those observed in cancer cells following treatment with splicing inhibitors (Table 1, Fig. 1F, Extended Data Fig. 1F, Supplementary Table S1-2). Furthermore, we scrutinized publicly available datasets of K562 and HepG2 cell lines after knockdown of 68 and 70 different splicing factors, respectively, from the ENCODE project, identifying similar inhibitor-induced transcriptomic changes alongside knockdown of other subunits of the U2 snRNP complex (Fig. 1G).

### Pharmacological inhibition of SF3B1 causes a global decrease in the abundance of DNA repair proteins, resulting in DNA damage

To be able to reduce therapeutic doses of splicing inhibitors, we examined whether low dose of splicing inhibitor could produce effects in tumor cells similar to those observed above. We conducted RNA-seq analysis of ovarian cancer cell line SKOV3 24 and 48 hours after treatment with a low dose (1.56 nM, IC30) of splicing inhibitor pladienolide B (Pl-B) (Extended Data Fig. 2A-B). Treatment with this low dose of Pl-B resulted in altered gene expression and pre-mRNA splicing, mirroring what we observed in our meta-analysis of various cancer cell lines after treatment with the Pl-B derivative, E7107. Cells treated with low dose of Pl-B also exhibited decreased expression of genes involved in DNA repair and replication (Fig. 2A, Supplementary Table S2), as well as extensive exon skipping estimated to involve approximately 11,000 events (Extended Data Fig. 2C, Supplementary Table S1). Recognizing the limitations of RNA-seq in accurately reflecting changes in protein abundance, we complemented our analysis with LC-MS/MS based proteomic profiling of SKOV3 cells 24 and 48 hours after Pl-B treatment (Extended Data Fig. 2D, Supplementary Table S3). The alterations observed in this proteomic data aligned with the findings reported in our RNA-seq analysis (Fig. 2A).

**Figure 2.**
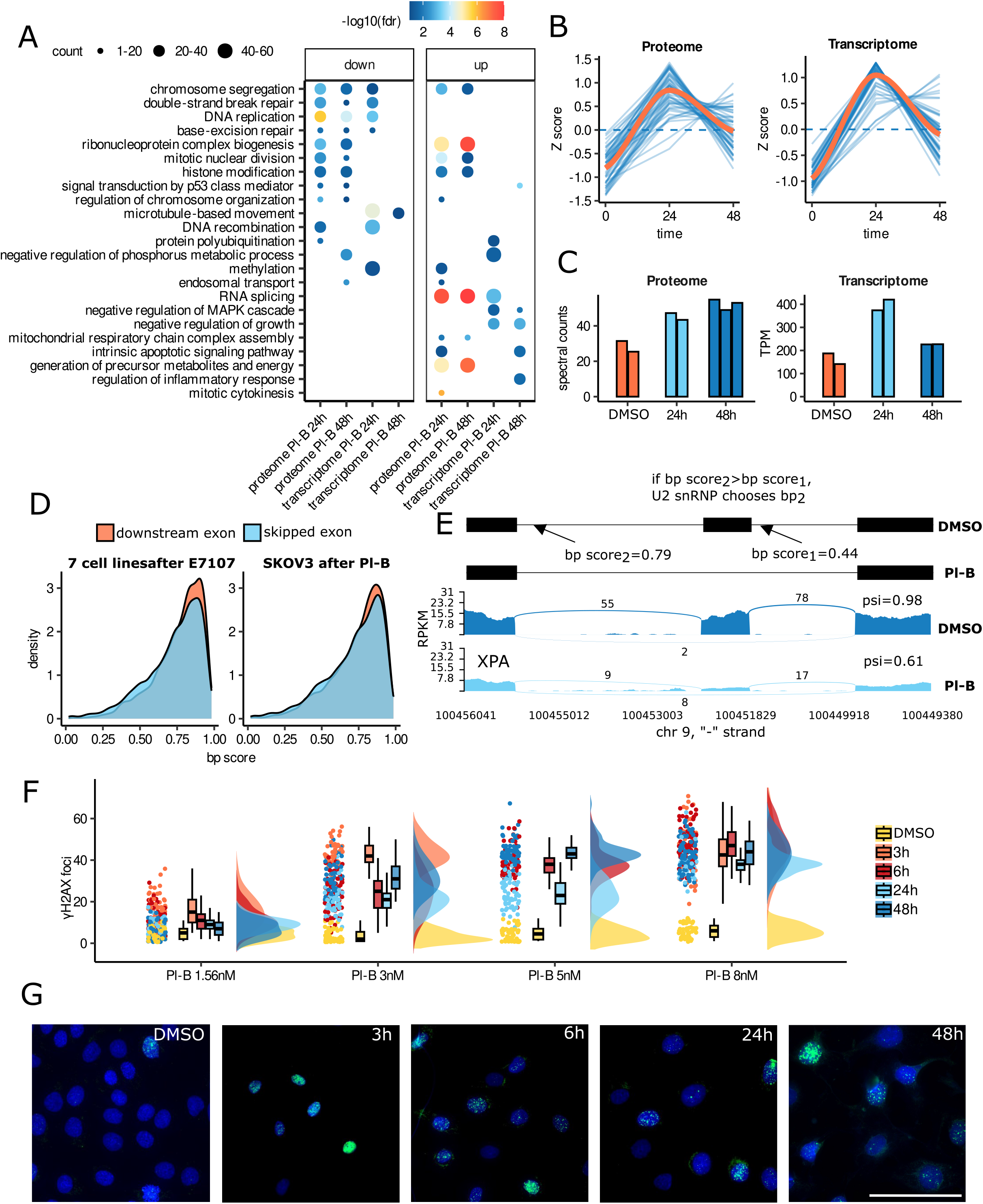
A rise in the expression of splicing factors leads to exon skipping after pladienolide B treatment. **A** – Dot plot illustrating Reactome pathway enrichment analyses (FDR < 0.05) of differentially expressed genes/proteins in SKOV3 cells 24 and 48 hours after pladienolide B (1.56 nM) treatment. Dot size corresponds to the count of genes/proteins enriched in the pathway, and color indicates pathway enrichment significance. **B** – Time clusterization of spliceosomal protein (left panel) and gene (right panel) expression in SKOV3 cells after pladienolide B treatment. Displayed are genes/proteins that significantly changed expression 24 hours after pladienolide B treatment. Y-axis represents mean-centered log2 (TPM+1) value or mean-centered log2 (spectral count+1) value. Blue denotes expression levels of each spliceosomal gene/protein, while orange represents their mean expression. **C** – Bar plots showing TPM or spectral count values of SF3B1 after pladienolide B (1.56 nM) treatment of SKOV3 cells based on RNA-seq and proteomic data, respectively. **D** – Distribution of branch point signal scores around skipped exons and downstream exons, calculated by the Branchpointer tool. Paired Wilcoxon test was performed for branch point score comparison. **E** – Sashimi plots for the DNA repair gene *XPA* in SKOV3 cells untreated (dark blue) and treated with pladienolide B (light blue). The psi value indicates the splicing status of the exon. Branch point annotation around the skipped exon of *XPA* gene was calculated using the Branchpointer tool. **F** – Raincloud plots depicting the number of γH2AX foci per nucleus in SKOV3 cells treated with various concentrations of pladienolide B at different time points. The number of γH2AX foci was calculated using ImageJ software with FindFoci plugins. 60-200 cells were analyzed in each sample. **G** – Representative immunofluorescence images of SKOV3 cells stained for γH2AX (Ser139, green) and with DAPI (blue) after treatment with 8 nM pladienolide B at different time points. Scale bar = 100 μm.

To explore whether tumor cells would endeavor to compensate for the loss of SF3B1 activity under the influence of Pl-B, we monitored changes in gene expression and protein abundance across different time points. Indeed, 24 hours after Pl-B exposure, approximately 50 genes and proteins involved in the regulation of pre-mRNA splicing, including SF3B1, were significantly upregulated (Fig. 2B-C). Notably, 48 hours after Pl-B treatment, the cells were able to recover from this exposure and restore the expression levels of pre-mRNA splicing genes/proteins to baseline (Fig. 2B). Therefore, we hypothesized that such a compensatory response of cancer cells could lead to disruption in the coordinated splicing network, provoking the binding of the U2 snRNP complex to more conservative and stable branch point sequences, ultimately resulting in massive exon skipping events. To test this hypothesis, we analyzed branch point sequence elements near skipped exons provoked by splicing inhibitors. Results revealed that splicing inhibitors caused skipping of exons characterized by significantly weakened branch points, particularly exhibiting more variation than upstream exons (Fig. 2D-E).

According to our RNA-seq data analysis, pharmacological inhibition of SF3B1 results in a decreased expression of crucial DNA repair genes, including *RAD51*, *RAD52*, *TOP2A*, *TOP2B*, and *CHEK2*. Inhibition of these genes have been previously shown to induce DNA breaks^27–30^. To investigate whether the splicing inhibitor leads to DNA damage, we assessed the phosphorylation level of histone H2AX (γH2AX) as an early marker of DNA damage response^31^. SKOV3 cells were subjected to γH2AX foci immunofluorescence staining at various time points (3, 6, 24, or 48 hours) after treatment with different doses of Pl-B (1.56-8 nM) (Fig. 2F-G). Our findings revealed that a low dose of Pl-B (1.56 nM) increases the number of γH2AX foci only at early time intervals (3-6 hours), while high doses of Pl-B (3-8 nM) cause a significant increase in γH2AX foci for a longer period (3-48 hours), indicating that pharmacological inhibition of SF3B1 leads to DNA damage. Collectively, these data show that splicing inhibitors could impair the DNA damage repair pathway in cancer cells, hence, targeting this vulnerability could increase therapeutic response.

### Impaired NMD functionality upon exposure to high splicing inhibitor doses

In this investigation, a robust correlation emerged between genes downregulated in response to splicing inhibitors and those exhibiting significantly altered exon skipping events. Consequently, we sought to ascertain whether these splicing events triggered the nonsense-mediated decay (NMD) process. For this purpose, we conducted an integrated bioinformatic analysis on seven publicly available RNA-seq datasets generated from human cells with knockdown of various NMD factors, including *UPF1*, *SMG6*, or *XRN1*, to compile a list of splicing events that could potentially serve as NMD targets (Table 1). Our analysis demonstrated a significant overlap between exon skipping events induced by E7107 or Pl-B treatment and exons that were skipped in mRNAs accumulated after the knockdown of core NMD factors (Fig. 3A-B). Functional enrichment analysis of the overlapping genes identified the “DNA Repair” pathway as the most significantly enriched (Fig. 3A-B). To validate our findings from alternative splicing analysis, we selected several DNA repair genes (*XPA*, *RAD51*, *CHEK2*, and *RIF1*) to conduct semi-quantitative RT-PCR, selecting two events for each gene based on whether they were undergoing NMD degradation or not. The RT-PCR results for all eight alternative splicing events tested on these four genes were consistent with our RNA-seq data (Fig. 3C, Extended Data Fig. 3A). Furthermore, to assess transcript stability and the level of NMD degradation, we correlated the results of nascent RNA-seq data (TT-seq) with paired bulk RNA-seq data analyses of CUTLL1 lymphoma cells after treatment with E7107 (Table 1). We observed a more substantial downregulation of DNA repair genes in RNA-seq data compared to the paired TT-seq data (Fig. 3D-E, Extended Data Fig. 3B), further confirming our hypothesis that downregulation of DNA repair genes is likely associated with their mRNA degradation.

**Figure 3.**
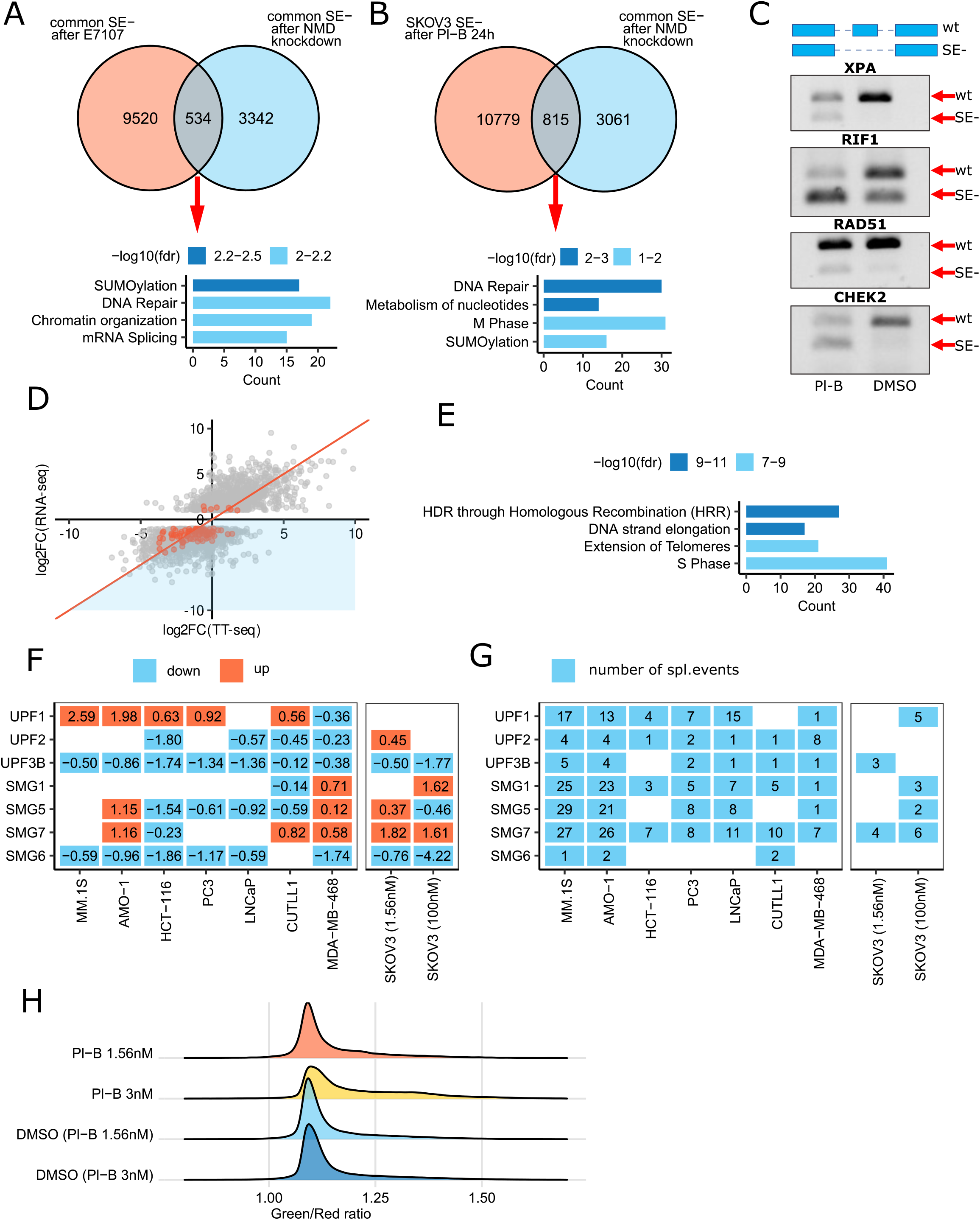
Impaired NMD functionality upon exposure to high splicing inhibitor doses. **A** – Venn diagram showcasing the overlap between skipped exons (SE) shared across 7 cancer cell lines after E7107 treatment and those present in datasets following NMD complex subunit knockdown. Reactome pathway enrichment analysis of overlapping genes is displayed below. The blue color scale indicates FDR values. **B** – Venn diagram depicting the overlap between exons skipped in SKOV3 cells treated with pladienolide B (1.56 nM) for 24 hours and those common in datasets after NMD complex subunit knockdown. Reactome pathway enrichment analysis of overlapping genes is presented below. The blue color scale indicates FDR values. **С** – RT-PCR results for the validation of exon skipping events in *XPA*, *RAD51*, *RIF1*, and *CHEK2* mRNAs in SKOV3 cells treated with 1.56 nM pladienolide B for 24 hours. “wt” signifies the wild- type transcript, “SE-” indicates the transcript with an exon skipping event. **D** – Scatter plot illustrating the differential gene expression (log2-fold change) of nascent RNAs from TT-seq analysis (x-axis) and mRNAs from RNA-seq analysis (y-axis) for CUTLL1 cells treated with E7107 for 24 hours compared to untreated CUTLL1 cells. Orange color highlights differentially expressed DNA repair genes. **E** – Functional enrichment analysis of differentially downregulated genes (FDR < 0.05 and log2- fold change < −1) with lower expression in RNA-seq data compared to TT-seq data. **F** – Heat map represents the log2-fold change values of main NMD factors (*UPF1, UPF2, UPF3B, SMG1, SMG5, SMG7, SMG6*) expression after treatment of different cancer cell lines with splicing inhibitors. Downregulated genes are marked in blue, upregulated genes are marked in orange (FDR < 0.05). The x-axis shows the name of the cancer cell line; the y-axis shows gene names. **G** – Heat map indicating the number of differential splicing events identified in mRNAs of the NMD factors (*UPF1, UPF2, UPF3B, SMG1, SMG5, SMG7, SMG6*) after treatment of different cancer cell lines with splicing inhibitors. The x-axis shows the name of the cancer cell line; the y- axis shows gene names. **H** – Representative density plots illustrating NMD activity determined by fluorescence reporter system analysis of SKOV3 cells treated with different concentrations of Pl-B. As a control, cells were treated with volumes of DMSO equivalent to Pl-B. The x-axis shows the ratio of fluorescence intensities of two proteins: one (green) is encoded by the NMD-targeted transcript, and the other (red) serves as an expression efficiency control. A higher ratio of green/red signals indicates lower NMD activity.

Simultaneously, we noted extensive splicing perturbation among transcripts related to key NMD factors (*UPF1, UPF2, UPF3B, SMG1, SMG5, SMG7, SMG6*) across various cancer cell lines after exposure to splicing inhibitors (Fig. 3F-G). Previous studies have demonstrated that NMD degradation can occur through two independent pathways^32^, regulated by either SMG5 and SMG7 proteins or SMG6 protein. Here, we observed a considerable decrease in SMG6 expression after treatment with splicing inhibitors across several tumor cell lines used in our study (Fig. 3F). Additionally, our RNA-seq of SKOV3 cells exposed to a high dose (100 nM) of Pl-B revealed an even greater reduction in SMG6 expression (18-fold decrease) compared to low-dose exposure (Fig. 3F). Based on these findings, we hypothesized that high doses of splicing inhibitors might cause a widespread increase in nonsense-bearing transcripts by reducing the activity of the NMD machinery. To test this hypothesis, we measured the activity of the NMD pathway in response to different concentrations of Pl-B using a reporter system^33^ that consisted of two plasmids: the first one encoded mRNA for the green fluorescent protein TagGFP2, which was NMD-targeted due to the presence of a premature termination codon, while the second encoded mRNA for the red fluorescent protein Katushka, which was NMD-insensitive. Our results showed that a higher Pl-B dose (3 nM; IC45) significantly reduced the NMD machinery’s activity, while at a low dose (1.56 nM; IC30), NMD activity remained largely unaffected, consistent with our RNA-seq data analysis, thus supporting our hypothesis (Fig. 3H, Extended Data Fig. 3C).

### Novel therapeutic approaches exploiting DNA repair vulnerability induced by splicing inhibitors

In our analysis of RNA-seq data of eight tumor cell lines after exposure to splicing inhibitors, a consistent decrease in the expression of 94 DNA damage repair genes was observed in at least half of the cell lines, irrespective of tumor type (Fig. 4A, Supplementary Table S4).

**Figure 4.**
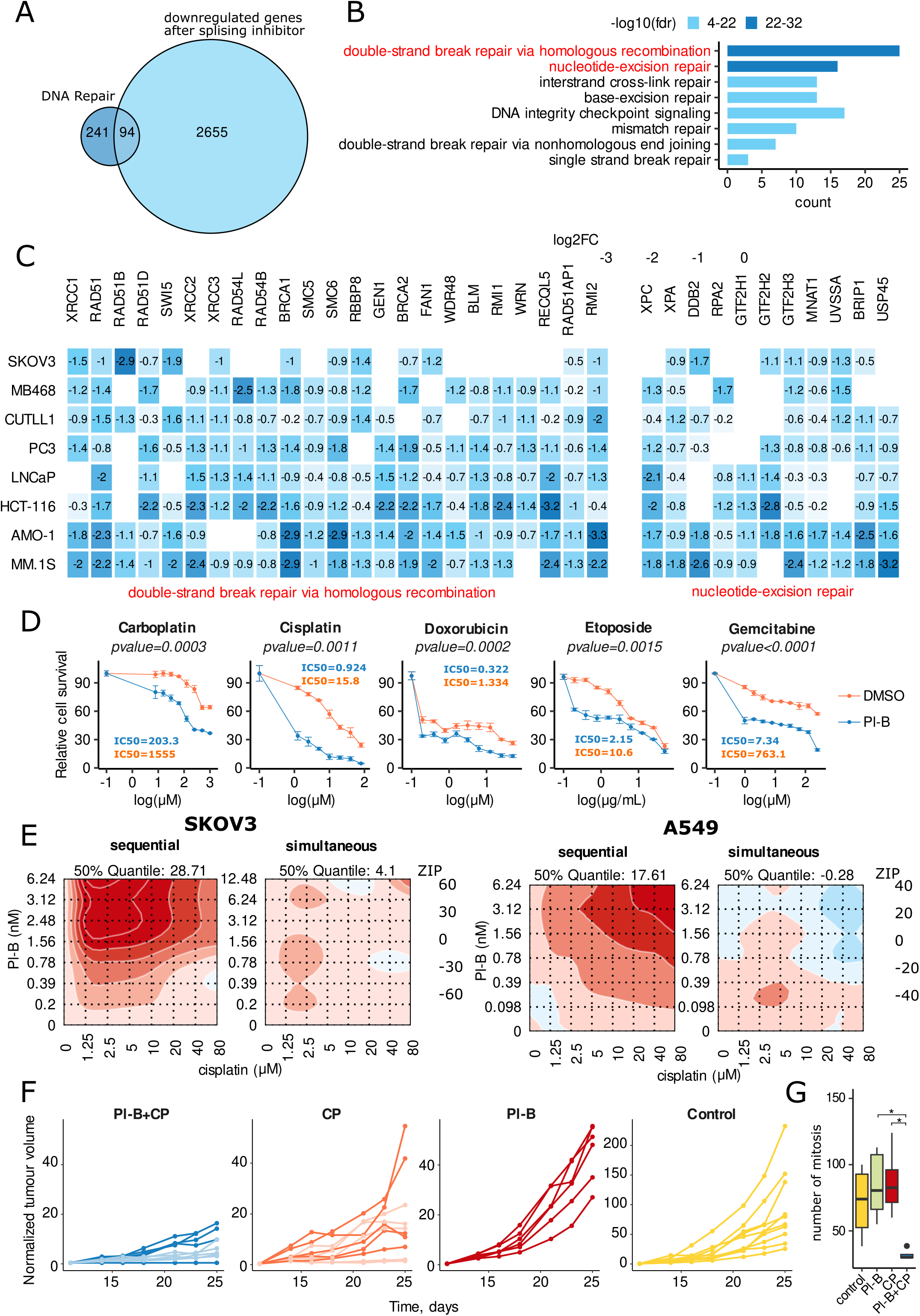
Novel therapeutic approaches exploiting DNA repair vulnerability induced by splicing inhibitors. **A** – Venn diagram showcasing the overlap between commonly downregulated genes in 8 cancer cell lines after treatment with splicing inhibitors and the list of DNA repair genes (according to the Reactome database and MD Anderson Cancer Center). A gene was considered “common” if its expression decreased in at least half of cell lines with FDR < 0.05 and log2-fold change < (−0.7). **B** – Functional annotation analysis of common DNA repair genes downregulated in 8 cancer cell lines after splicing inhibitor treatment. Eight GO terms, comprehensively representing known DNA repair mechanisms, were selected for analysis. The blue color scale indicates FDR values. **C** – Heat map representing log2-fold change values for differentially expressed genes in 8 cancer cell lines after splicing inhibitor treatment. The graph includes genes in GO terms “double-strand break repair via homologous recombination” or “nucleotide excision repair,” found to be differentially expressed in at least half of the cancer cell lines. The graph labels log2FC only for genes with FDR < 0.05. The blue color scale indicates log2-fold change values. **D** – Dose-response curves obtained by MTT assay of SKOV3 cells pretreated with 1.56 nM pladienolide B (48 hours) followed by treatment with different concentrations of carboplatin, cisplatin, doxorubicin, etoposide, or gemcitabine (48 hours). Each data point represents mean values ± SD (n = 3). IC50 values were determined using GraphPad Prism software (normalized model to data with nonlinear regression). Significance determined using a paired, two-tailed Student’s t-test, p-values presented on the image. **E** – Synergy landscapes for SKOV3 and A549 cells after simultaneous or sequential treatment with different doses of pladienolide B (y-axis) and cisplatin (x-axis). Zero-Interaction Potency (ZIP) synergy scores calculated for the combination of pladienolide B (0 nM–6.24 nM) and cisplatin (0 µM–80 µM). For simultaneous regimen, сells were treated with two drugs simultaneously (48 hours). For sequential regimen, cells were pretreated with different concentrations of pladienolide B (48 hours) followed by treatment with different concentrations of cisplatin for 48 hours. Cell survival was calculated in comparison to untreated cells (DMSO only). ZIP values lower than 0 indicate an antagonistic effect of the drug combination (blue), 0 - 10 indicate an additive effect (from white to light red), values above 10 (corresponding to a deviation from the reference model above 10%) indicate synergy (dark red). **F** – CT26 tumor progression in mice for each *in vivo* treatment group. Normalized tumor volume (NTV) calculated as (Vx - V1), where Vx is tumor volume on day X, and V1 is tumor volume at the start of treatment (blue - PL+CP5; light blue - PL+CP2.5; orange - CP5; light orange - CP2.5). “Pl-B + CP5” - group of mice treated with combination of pladienolide B (1 mg/kg) and cisplatin (5mg/kg) (n=5), “Pl-B + CP2.5” - group of mice treated with combination of pladienolide B (1 mg/kg) and cisplatin (2.5mg/kg) (n=5), “CP5” - group of mice treated with cisplatin (5mg/kg) (n=5), “CP2.5” - group of mice treated with cisplatin (2.5mg/kg) (n=5), “Pl-B” - group of mice treated with pladienolide B (1 mg/kg) (n=6), “control” - control group of untreated mice (n=10). **G** – Box plot represents the number of mitoses in CT26 tumors for each *ex vivo* treatment group. Number of mitoses calculated using QuPatch and ImageJ software. Asterisks denote statistically significant differences determined using Wilcoxon test: * means p < 0.05.

Notably, these DNA damage repair genes are implicated in homologous recombination and the nucleotide excision repair pathway (Fig. 4B-C). Given previous findings that the knockout of even one of these DNA damage response genes can enhance the cytotoxicity of tumor cells to genotoxic agents^34^, we hypothesized that a low dose of a splicing inhibitor, in combination with a DNA-damaging agent, could yield a synergistic effect. To test this hypothesis, we assessed the efficacy of the combined effect of a low dose of the splicing inhibitor Pl-B and various clinically used DNA-damaging agents (carboplatin, cisplatin, doxorubicin, etoposide, gemcitabine) in a simultaneous or sequential regimen on the ovarian adenocarcinoma cell line SKOV3 (Fig. 4D, Extended Data Fig. 4A). Significantly, only the sequential regimen of drug administration markedly increased the sensitivity of SKOV3 cells to all tested DNA-damaging agents (Fig. 4D, Extended Data Fig. 4A). The combination of Pl-B and cisplatin was selected for further experiments, given cisplatin’s widespread use as a chemotherapeutic drug^35^. To optimize the incubation time with Pl-B for a stronger response of cancer cells to cisplatin, we conducted time-dependent pre-incubation experiments, revealing that the greatest increase in sensitivity occurred at 48 hours after Pl-B addition, where the IC50 value of cisplatin decreased by 11 times (Extended Data Fig. 4B). Similar results were obtained for cisplatin and another splicing inhibitor, H3B-8800, which is under clinical trials (Extended Data Fig. 5H, I). This result was validated by detecting the level of apoptosis using flow cytometry (Extended Data Fig. 4C). The proposed drug combination in a sequential regimen was also shown to be effective in other cell lines representing different cancer types (Extended Data Fig. 4D, Extended Data Fig. 5A). Intriguingly, pre-incubation of human primary fibroblasts and fallopian tube secretory epithelia cells (FT282) with Pl-B did not increase their sensitivity to cisplatin (Extended Data Fig. 5B-C), indicating that this therapeutic vulnerability might be exclusive to cancer cells, underscoring the potential clinical relevance of our findings. Even Pl-B-resistant (Extended Data Fig. 5D) cancer cells adapted to suppressed SF3B1 function exhibited increased sensitivity when treated with cisplatin (Extended Data Fig. 5E). Additionally, cisplatin-resistant cancer cells (Extended Data Fig. 5F) became sensitive to cisplatin after pre-incubation with Pl-B (Extended Data Fig. 5G).

We further evaluated the synergy score between Pl-B and cisplatin in several cancer cell lines representing different tumor types (ovarian cancer SKOV3, non-small cell lung cancer A549, human hepatocyte carcinoma HepG2, and colorectal cancer HT29) using Zero-Interaction Potency (ZIP) analysis. Results demonstrated a high synergy score exclusively in the sequential regimen at nanomolar concentrations of Pl-B (Fig. 4E, Extended Data Fig. 5J-M). There was a 6-10 fold increase in the effective potency of cisplatin during sequential therapy, indicating a true synergistic effect rather than additive drug compounding.

To assess whether splicing inhibitors sensitize malignant tumors to DNA-damaging agents *in vivo*, we established a subcutaneous murine syngeneic tumor model CT26 expressing the far-red fluorescent protein eqFP650 (Extended Data Fig. 6C-F). We first confirmed the efficacy of the proposed drug combination on this cell line in a sequential regimen *in vitro* (Extended Data Fig. 6A-B). Subsequently, we demonstrated that treatment of mice with cisplatin or a combination of cisplatin and Pl-B significantly reduced tumor burden compared to the untreated control group (Fig. 4F). Notably, the most significant trend in the decline of tumor growth rate *in vivo* was observed for tumors treated with a combination of Pl-B and cisplatin. It is worth noting that tumor growth was reduced in all tumors in the group treated with a combination of Pl-B and cisplatin at different doses of cisplatin (Fig. 4F; Extended Data Fig. 6D-F). In contrast, when tumors were treated with cisplatin alone, we did not observe a therapeutic effect in 50% of the tumors (Fig. 4F). Histopathological analysis of treated tumor tissues revealed cell death (karyopyknosis, karyorrhexis, and apoptotic bodies), cell shrinkage, nuclear damage, and notable nuclear polymorphism. Tumor tissues treated with a combination of Pl-B and cisplatin exhibited the most pronounced nuclear polymorphism, leading to the formation of tumors with undefined and irregular morphology (Extended Data Fig. 6G). As counting mitotic figures is considered one of the most important methods^36,37^ to assess an individual’s response to anticancer therapy, we counted single mitoses in tumors from different experimental animal groups. Tumors subjected to sequential therapy showed a significant decrease in the mitotic activity of cancer cells, predicting longer survival and a low probability of developing distant metastases^36,37^ (Fig. 4G).

### Splicing inhibitor pretreatment impairs activation of cellular pathways important for DNA damage response

To unravel the mechanistic insights behind the synergistic action of cisplatin and Pl-B, we conducted transcriptomic analysis of total mRNA extracted from SKOV3 cells exposed to Pl-B, cisplatin, or their combination, as illustrated in the scheme in Fig. 5A. Specifically, cells were pre-treated with low doses of Pl-B for 48 hours, followed by replacement of Pl-B-containing media with cisplatin-containing one for an additional 12 hours. As controls, SKOV3 cells were pre-incubated with either Pl-B or DMSO for 48 hours, followed by replacement with fresh medium without drugs, referred to as “Pl-B release” and “DMSO,” respectively. Although the low-dose Pl-B treatment induced significant changes in the expression profiles of cells (Fig. 2A-B), this change was reversible as expression profiles being restored to their previous values after 12 h release after Pl-B (Fig. 5B-C, Supplementary Table S5). Furthermore, principal component analysis demonstrated that SKOV3 cells pre-treated with Pl-B or DMSO activated distinct transcriptomic programs in response to cisplatin treatment (Fig. 5B). To identify the biological pathways contributing to the pronounced cytotoxicity of the proposed combination, we conducted a differential gene expression analysis. The Pl-B and cisplatin combination resulted in a significantly greater reduction in the expression of genes primarily involved in DNA repair, cell cycle regulation, SUMOylation, and regulation of TP53 activity compared to all other treatments (Fig. 5D). We also performed above-mentioned RNA-seq experiments on SKOV3 cells using the clinically relevant splicing inhibitor H3B-8800 instead of Pl-B (Table 1, Extended Data Fig. 7A, Supplementary Table S5). Combination treatment of H3B-8800 and cisplatin induced changes in gene expression in tumor cells similar to groups treated with the combination of Pl-B and cisplatin (Extended Data Fig. 7A-C, Supplementary Table S5). To determine whether the changes in transcriptome profiles observed in cancer cells with drug combination treatment were mediated by the prior action of a splicing inhibitor, we conducted clustering analysis of gene expression changes in SKOV3 cells after Pl-B treatment alone. Genes were categorized into two clusters: downregulated (cluster 1) and those without changes in expression (cluster 2) under the influence of Pl-B (Fig. 5E). Functional annotation of genes from cluster 1 revealed their predominant involvement in DNA repair (Fig. 5F). Meanwhile, the contribution of the drug combination (cluster 2) resulted in the downregulation of signaling pathways, such as cell cycle regulation, SUMOylation, and the regulation of TP53 activity (Fig. 5E). Thus, these data suggest that Pl-B reduces the expression of genes crucial for an effective response of cells to DNA damage insults, rendering them more susceptible to DNA-damaging drugs.

**Figure 5.**
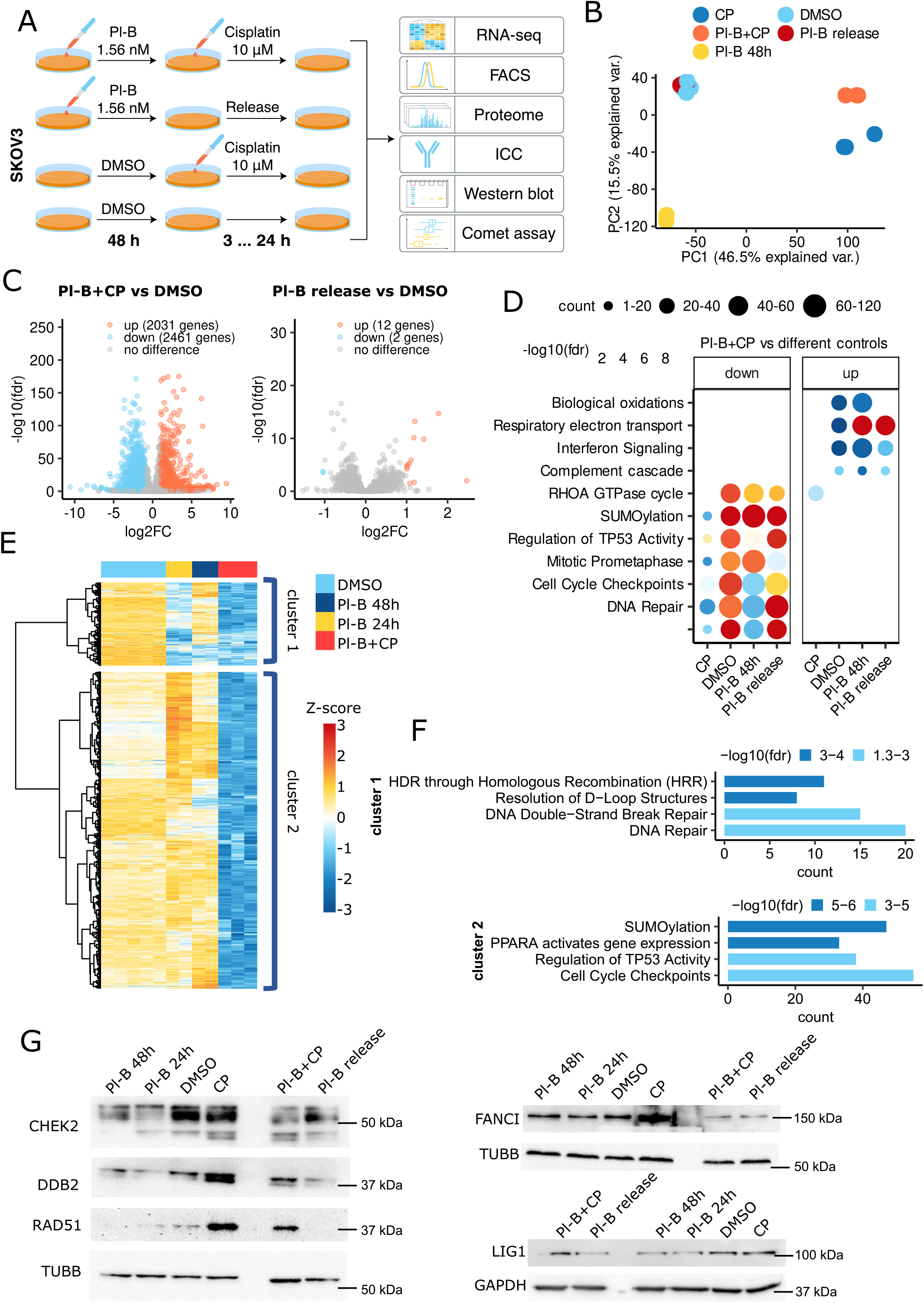
Splicing inhibitor pretreatment impairs activation of pathways important for cell response to DNA damage. **A** – Experimental workflow used to study differences in SKOV3 cells after treatment with pladienolide B (1.56 nM), cisplatin (10 µM) or their combination. DMSO-treated cells were taken as a control. ICC - immunocytochemistry. **B** – Principal component analysis (PCA) of the log2-transformed normalized gene expression values between control (DMSO) and treated samples. Each point represents the expression profile of one sample. Types of treatment: “DMSO” - cells treated with DMSO for 24 hours, “CP” - cells treated with cisplatin (10 µM) for 12 hours, “Pl-B 48h” - cells treated with pladienolide B (1.56 nM) for 48 hours, “Pl-B + CP” - cells pretreated with pladienolide B (1.56 nM) for 48 hours followed by cisplatin (10 µM) treatment for 12 hours, “Pl-B release” - cells treated with pladienolide B (1.56 nM) for 48 hours and then cultivated in fresh medium without a drug for 12 hours. **C** – Volcano plots highlight genes identified as differentially expressed (FDR < 0.1 and |log2-fold change| > 1) in SKOV3 cells after treatment with the combination of pladienolide B and cisplatin (Pl-B + CP) or pladienolide B (1.56 nM) for 48 hours and its removal from the medium (Pl-B release) compared to control DMSO-treated cells. **D** – Dot plot shows the Reactome pathways (FDR < 0.05) enrichment analysis of differentially expressed genes/proteins in SKOV3 cells treated with the proposed drug combination compared to different controls as described in Figure 5A. Dot size is based on gene/protein count enriched in the pathway, and dot color indicates pathway enrichment significance. **E** – Heat map represents RNA-Seq expression z-scores computed for genes that are differentially expressed (FDR < 0.1, log2-fold change < −1) between SKOV3 cells after treatment with DMSO or the drug combination. kMeans clustering was used for expression pattern division. **F** – Reactome Pathway enrichment analysis of two gene clusters from Figure 5E. Colors indicates the −log10(fdr) values. **G** – Western blot analysis of total lysates of SKOV3 cells under different types of treatment: “DMSO” - cells treated with DMSO for 24 hours, “CP” - cells treated with cisplatin (10 µM) for 12 hours, “Pl-B 24h” and “Pl-B 48h” - cells treated with pladienolide B (1.56 nM) for 24 and 48 hours, “Pl-B + CP” - cells pretreated with pladienolide B (1.56 nM) for 48 hours followed by cisplatin (10 µM) treatment for 12 hours, “Pl-B + release” - cells treated with pladienolide B (1.56 nM) for 48 hours and then cultivated in fresh medium without a drug for 12 hours.

Finally, we conducted LC-MS/MS analysis of cells treated with cisplatin alone or the combination of cisplatin and a splicing inhibitor (Pl-B or H3B-8800) to examine the proteomic changes in cancer cells 24 hours after treatment (Extended Data Fig. 7D-E, Supplementary Table S6). Combination treatment led to a significant downregulation of pathways associated with cell cycle regulation and DNA repair, where approximately 100 proteins, including XPA, DDB2, RAD51, CHEK2, TDP1, and others, were altered. This result aligns with our transcriptomic data (Fig. 5D). We further validated our findings using western blot analysis (Fig. 5G).

### Replicative stress as a key reason for the cytotoxic effect of the proposed combination therapy

Our analysis of transcriptomic and proteomic data (Fig. 1D, 2A, 5D) revealed that Pl-B significantly reduces the expression of proteins involved in DNA damage response, potentially leading to more extensive DNA damage in the case of combining Pl-B with DNA-damaging agents. To test this hypothesis, we measured DNA damage using comet assays and analyzed the DNA repair dynamics by assessing the phosphorylation levels of H2AX in SKOV3 cells after treatment with Pl-B, cisplatin, or their combination. We noticed a decrease in the number of γH2AX foci, while observing a greater number of DNA breaks in cells treated with the drug combination compared to cisplatin alone (Fig. 6A-C, Extended Data Fig. 8A-B). This effect may be attributed to the fact that Pl-B either disrupts the sensing of DNA breaks due to the suppression of the DNA repair process or overloads the DNA repair machinery due to excessive replicative stress, making cells more susceptible to subsequent cisplatin treatment. To confirm this hypothesis, we assessed the activation of three main phosphatidylinositol 3-kinase-related kinases, ATM, ATR, and DNA-PK, which are responsible for the phosphorylation of H2AX^38,39^. The drug combination treatment of SKOV3 cells reduced the phosphorylation levels of ATR, ATM, and DNA-PK compared to treatment with cisplatin alone, potentially blocking subsequent cascades of DNA repair reactions (Fig. 6D-F). Next, we investigated the induction of replication stress, analyzing the number of R-loops (toxic DNA:RNA hybrids)^40^ and the phosphorylation level of RPA2 (Ser33)^41^ in SKOV3 cells after different treatments. The fluorescence signals of R-loops and phosphoRPA2 significantly increased in cells 6 hours after treatment with the drug combination compared to cisplatin or Pl-B treatments alone (Fig. 6G-H, Extended Data Fig. 8C-D), indicating increased early replicative stress.

**Figure 6.**
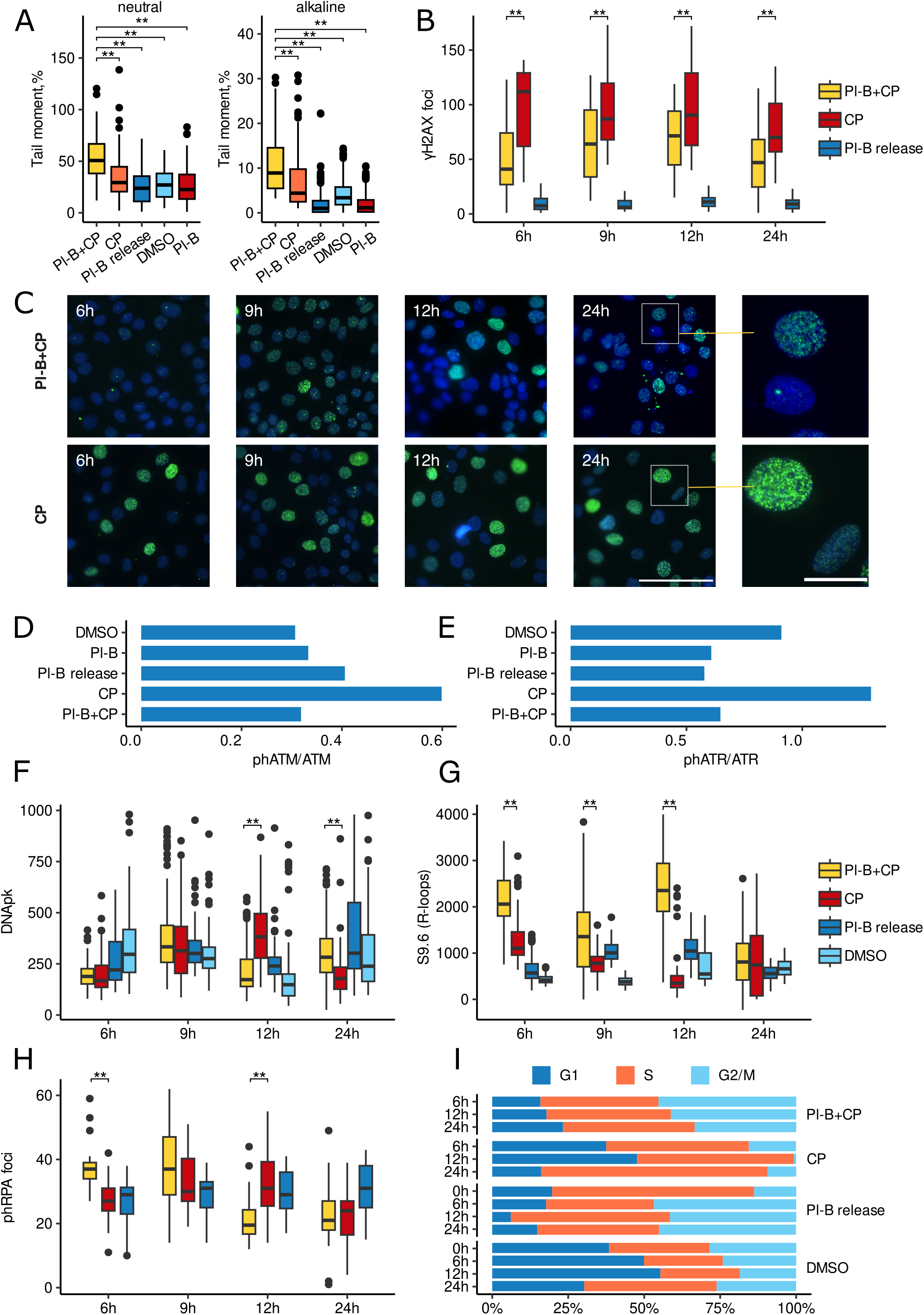
Replicative stress as a key reason for the cytotoxic effect of the proposed combination therapy. **A** – Box plots of tail moments from neutral (double-stranded breaks; on the left) and alkaline (single and double-stranded breaks; on the right)) comet assays of SKOV3 cells after different types of treatment. Tail moment was defined as the product of the tail length and the fraction of total DNA in the tail (Tail moment = tail length x % of DNA in the tail) and was quantified using the CometScore 2.0 software. 60-100 cells were analyzed in each sample. Experiments were performed in triplicate. Asterisks denote statistically significant differences determined using a paired, two-tailed Student’s t-test: * means p < 0.05; ** means p < 0.01. **B** – Box plots show the number of γH2AX foci per nucleus in SKOV3 cells under different types of treatment. The number of γH2AX foci was calculated using ImageJ software with FindFoci plugins. 60-200 cells were analyzed in each sample. **С** – Representative immunofluorescence images of SKOV3 cells stained for phosphorylated H2AX (Ser139, green) and with DAPI (blue) after various treatments. Scale bar = 100 μm and 25 μm. **D** – Bar graph demonstrates the ratio of levels of phosphorylated and total ATM forms according to immunofluorescence staining followed by flow cytometry analysis of SKOV3 cells under different types of treatment. **E** – Bar graph demonstrates the ratio of levels of phosphorylated and total ATR forms according to immunofluorescence staining followed by flow cytometry analysis of SKOV3 cells under different types of treatment. **F** – Box plots show the fluorescence intensity of DNApk in SKOV3 cells under different types of treatment. The fluorescence intensity of DNApk was measured using ImageJ software. 60-200 cells were analyzed in each sample. **G** – Box plots show the fluorescence intensity of R-loops in SKOV3 cells under different types of treatment. The fluorescence intensity of R-loops was measured using ImageJ software. 80-200 cells were analyzed in each sample. **H** – Box plots show the number of phosphorylated RPA2 foci per nucleus in SKOV3 cells under different types of treatment. The number of phosphorylated RPA2 foci was calculated using ImageJ software with FindFoci plugins. 60-200 cells were analyzed in each sample. **I** – Cell cycle analysis with flow cytometry of SKOV3 cells stained with DAPI under different types of treatment. Stacked bar graphs show the percentage of cells in different phases of the cell cycle. Percentage of cells in G1, S, and G2/M phases was calculated with NovoExpress software. “Pl-B + CP” - SKOV3 cells were pretreated with 1.56 nM of pladienolide B for 48 hours (in A, D, E) or 6h, 9h, 12h and 24h (in B, C, F, G, H, I), then the medium was changed to the medium with 10 µM of cisplatin for 24 hours; “CP” - SKOV3 cells were incubated with DMSO for 48 hours (in A, D, E) or 6h, 9h, 12h and 24h (in B, C, F, G, H, I), then the medium was changed to the medium with 10 µM of cisplatin for 24 hours (in A, D, E) or 6h, 9h, 12h and 24h (in B, C, F, G, H, I); “Pl-B release” - SKOV3 cells were incubated with 1.56 nM of pladienolide B for 48 hours, then the medium was changed to fresh one without pladienolide B for 24 hours (in A, D, E) or 6h, 9h, 12h and 24h (in B, F, G, H, I); “DMSO” - SKOV3 cells were incubated with DMSO for 48 hours (in A, D, E) or 6h, 9h, 12h and 24h (in F, G); “Pl-B” - SKOV3 cells were treated with 1.56 nM of pladienolide B for 48 hours (in A, D, E). Asterisks denote statistically significant differences determined using Wilcoxon test: ** means p < 0.01 (in B, F, G, H).

While analyzing γH2AX staining, we observed differences in the patterns of staining between cell populations after drug treatments. Pannuclear staining of γH2AX was more pronounced in SKOV3 cells after cisplatin treatment, which is typical for S-phase cells, while separate large foci appeared presumably in G1- and G2-phases of the cell cycle^42^. Based on this result and our omics data, we speculated that the drug combination prevents cancer cells from arresting in the S phase, required for proper DNA repair. Indeed, cell cycle analysis showed that adding cisplatin to cells after pre-incubation with Pl-B did not significantly accumulate them at the S phase^43^ (Fig. 6I). Thus, the proposed drug combination in sequential regimen provokes the activation of replication stress, probably due to the formation of an excessive number of toxic R-loops and unrepaired DNA lesions, leading to increased efficacy of drugs on cancer cells.

## DISCUSSION

Splicing inhibitors represent a promising class of anticancer drugs^6^, though their use as standalone therapies at therapeutic doses is challenging due to significant side effects^7^. Moreover, the precise mechanism by which splicing inhibitors induce cancer cell death remains elusive^44^, despite their incorporation into clinical trials. In this study, we conducted an extensive meta-analysis of the transcriptomic responses of diverse cancer cell lines to splicing inhibitors. Surprisingly, we observed consistent splicing changes across cell lines representing different cancer types, indicating a commonality in their response to these inhibitors. Contrary to our expectations, splicing events induced by splicing inhibitors, primarily targeting the SF3B1 protein, a key subunit of the U2 snRNP complex, were characterized by a substantial occurrence of exon skipping (60-70%), rather than the anticipated intron retention. This unexpected phenomenon aligns with prior reports on the splicing inhibitors E7107 and H3B-8800, highlighting the prevalence of exon skipping in leukemia cell lines^16,19^. Our investigation further revealed that large-scale splicing alterations could be attributed not only to SF3B1 inactivation but also to a compensatory response by the cell, involving increased expression of approximately 50 spliceosomal proteins, including SF3B1 itself. This compensatory mechanism, previously observed in splicing factors of the hnRNP or SR families, involves negative autoregulatory pathways^45–47^. These splicing factors are known to participate in the splicing of their own pre-mRNA generating unproductive isoforms with premature termination codons, which are then degraded by the NMD pathway^24^. However, the mechanism by which SF3B1 initiates a negative autoregulation process remains unclear.

The consequential exon skipping events lead to the generation of unproductive transcripts harboring premature termination codons in genes responsible for DNA repair factors. These transcripts undergo degradation through the NMD pathway, reducing the abundance of corresponding proteins and potentially impairing the DNA repair process. However, our analysis of paired TT-seq and RNA-seq data unveiled a global decrease in gene expression, indicating that some genes were downregulated for reasons unrelated to NMD degradation. This observation is consistent with previous studies demonstrating a reduction in global RNA synthesis, attributed to the splicing inhibitors’ ability to diminish RNA polymerase II elongation velocity^18,48^.

Notably, we demonstrated that knockdown of SF3B1 or treatment with splicing inhibitors, rather than mutations in SF3B1, resulted in significant splicing dysregulation of DNA repair genes. Based on these findings, we propose that a combined approach employing splicing inhibitors along with clinically used DNA-damaging agents may offer enhanced efficacy in clinical practice.

Current clinical trials predominantly explore splicing inhibitors as single agents^7–10^. However, our study suggests that their combination with DNA-damaging agents may yield better outcomes. Notably, we observed the optimal time window between administration of splicing inhibitor and the second DNA-damaging agent underscoring the importance of precise treatment scheduling. Here we showed that cancer cells were able to recover after *in vitro* treatment with low doses of Pl-B and restore the expression levels of genes corresponding to untreated cells. These results are consistent with preclinical observations showed that maximum changes in pre-mRNA splicing of selected genes in patient samples were noticed at 4 - 10 hours after H3B-8800 dosing and then pre-mRNA splicing perturbations diminished^9^.

While previous research has proposed combining splicing inhibitors with various agents such as proteasome inhibitors, CHEK2 inhibitors, as well as with several DNA-damaging agents for treating leukemia or myeloma cells^16,19,49^, our study demonstrates that the proposed combination enhances sensitivity to DNA-damaging agents across different cancer cell lines, irrespective of tumor type or splicing factor mutation status. Importantly, this combination strategy exhibits minimal toxicity towards non-tumor cells, and the development of resistance to splicing inhibitors does not compromise sensitivity to the proposed combination.

Our findings also align with the emerging concept of a “BRCAness” phenotype^50,51^, a signature observed in cancers lacking mutations in BRCA1 or other DNA repair genes^51–54^. Splicing inhibitor pretreatment induced a BRCAness-like phenotype in different cancer cells, not only by downregulating DNA repair proteins but also by increasing replication stress which is a main reason inducing the cell death in the case of BRCA-deficient tumors^54,55^. We hypothesized that the replication stress, observed in tumor cells under the influence of the proposed combination, arises from the cells’ inability to effectively activate DNA repair cascades, leading to unrepaired DNA lesions that act as barriers to replication fork progression^56^. Our study presents a compelling case for the application of combined splicing inhibitor and DNA-damaging agent therapy in the treatment of various tumors, promising improved efficacy and minimal toxicity.

## Acknowledgments

This work was supported by grant 075-15-2019-1669 from the Ministry of Science and Higher Education of the Russian Federation (K.S.A., M.A.L., P.V.S., O.M.I., K.M.K.) for RNAseq analyses); by the Russian Science Foundation project no. 22-15-00462 (K.S.A., V.O.S., M.M.L., A.N.K., P.V.S., O.M.I., G.P.A.) for proteomic analysis, bioinformatics and experiments with cell cultures). We thank Dr. Sonya Bastola for the critical reading and editing of the manuscript.

## Author contributions

Conceptualization: K.S.A., M.M.L., and V.O.S.; Data curation: K.S.A., M.M.L., and V.O.S.; Formal analysis: K.S.A., M.M.L., G.P.A., and V.O.S.; Funding acquisition: K.S.A. and V.O.S.; Investigation: V.O.S., K.S.A., M.M.L., P.V.S., G.P.A., M.S.P., O.M.I., K.M.K., V.A.V., A.V.L., A.K.V., E.N.M., A.V.K., E.A.V., R.V.D., A.N.T., M.A.L., O.L.K., M.P.N., V.S.B., Zh.L., Z.W. and V.M.G.; Methodology: V.O.S., K.S.A., M.M.L., P.V.S., G.P.A., A.M.E., M.S.P., O.M.I., R.H.Z., K.M.K., V.A.V., N.M.M., A.V.L., A.K.V., E.N.M., R.V.D., V.S.B., and V.M.G.; Project administration: V.O.S.; Resources: V.O.S. and G.P.A.; Software: K.S.A., A.N.K. and G.P.A.; Supervision: K.S.A., V.M.G., and V.O.S.; Validation: M.S.P., K.S.A., G.P.A., and M.M.L.; Visualization: K.S.A., M.M.L., and A.N.K.; Writing – original draft: K.S.A., M.M.L., and V.O.S; Writing – review & editing: M.S.P., M.A.L., O.L.K., and V.M.G. All authors have read and agreed to the published version of the manuscript.

## Availability of data and materials

### Lead contact

Further information and requests for resources and reagents should be directed to and will be fulfilled by the Lead Contact, Victoria O. Shender.

### Data and code availability

All RNAseq data have been deposited at the Gene Expression Omnibus database (GEO) under the accession number — GSE227111— and are publicly available as of the date of publication. All proteomic datasets have been deposited to the ProteomeXchange Consortium via the PRIDE partner repository with the dataset identifier — PXD040874. All other data supporting the findings of this study are available from the corresponding author on reasonable request. This paper also analyzes existing, publicly available data. All accession numbers for these datasets are listed in Table 1. This paper does not report the original code. Any additional information required to reanalyze the data reported in this paper is available from the lead contact upon request.

### Competing interests

The authors declare the following competing financial interest(s): M.P.N. and E.N.M. are stakeholders of Abisense LLC which manufactures the LumoTrace FLUO bioimaging system.

## METHODS

### Alternative splicing analysis

Unprocessed RNA-Seq reads were trimmed using TrimGalore (version 0.6.6)^71^, wherein adapter/index sequences were trimmed, and reads shorter than 35 nucleotides were filtered. The trimmed RNA-Seq reads were then mapped to hg19 of Gencode (release 19) using STAR (v. 2.7.9a) with specified parameters from the STAR manual 2.7.0a. These parameters included: the maximum number of mismatches for paired reads was 4% of the read length, the maximum number of multiple alignments allowed for a read was 10, non-canonical junctions were removed, the minimum number of allowed splice overhangs was 8 for unannotated junctions and 1 for annotated junctions, the minimum intron length was 20, the maximum intron length was 1,000,000, and the number of “spurious” junctions was reduced. To compare splicing events in cancer cell lines after different treatments described in Table 1, rMATS(v.4.1.1)^72^ splicing tool was employed with recommended parameters (-c 0.0001 and -novel SS1). It has been repeatedly demonstrated that splicing events with low coverage lead to a low level of confidence in the psi value. To filter out low-confidence events, a custom script was written in the R programming language that calculates the number of reads covering exon-exons and exon-introns boundaries. If the average value of such reads among samples within one experiment was below 5, such splicing events were considered as noise events. The idea of this filtering was taken and adapted from the maser R/Bioconductor package. A splicing event was considered differential if the FDR was below 0.05 and the absolute value of dpsi exceeded 0.05. For the meta-analysis, a differential splicing event was considered “common” across different datasets if dpsi values were co-directed, splicing event coordinates were identical, and the event was deemed differential (FDR<0.05 |dpsi|>0.05) in at least half of the datasets.

Skipped exons and their upstream exons were analyzed using rtracklayer R/Bioconductor package^73^. Exon characteristics (sequences, rank in transcript, etc.) were obtained from Gencode annotation (release 19). Predicted splice branch point locations and their scores were downloaded from https://github.com/raphaelleman/BenchmarkBPprediction^74^. We collected branch point scores near skipped exons and downstream exons predicted by 5 different bioinformatic tools (SVM-BPfinder, BPP, Branchpointer, LaBranchoR and RNABPS). The paired Wilcoxon test was performed to compare the above described characteristics of the skipped exons and downstream exons; the differences were considered significant at fdr< 0.05.

### Differential gene expression analysis

The filtering and initial preparation of reads followed the same process as in the alternative splicing analysis. Subsequently, for each sample, transcript-level abundances were quantified using the quasi-mapping approach with Salmon (v. 0.12.0)^75^ on the hg19 Gencode reference transcriptome with default parameters. These transcript-level abundances were then aggregated to gene-level abundances using the tximport R/Bioconductor package (v. 1.16.1)^76^. Differential gene expression analysis was carried out with the DESeq2 R/Bioconductor package^77^ using the Wald test, applying an FDR cutoff of 0.05 and considering log2 fold change greater than the absolute value 1 for significance testing. Importantly, all the preceding steps were conducted separately for each dataset (Table 1). For the meta-analysis, a gene expression change was regarded as “common” across different datasets if two conditions were met: 1) log2 fold change values were co-directed, and 2) the gene expression was considered differential (FDR<0.05, |log2FC|>1) in at least half of the datasets.

### Functional enrichment analysis

Functional enrichment analysis for Reactome pathways and Gene Ontology (GO) terms was conducted using the clusterProfiler R/Bioconductor package, ReactomePA R/Bioconductor package, and topGO R/Bioconductor package. FDR cutoff of 0.05 was applied to determine statistical significance. The list of DNA repair genes utilized in this analysis was obtained from the Reactome database and www.mdanderson.org/documents/Labs/Wood-Laboratory/human-dna-repair-genes.html.

### TT-seq data analysis

The unprocessed reads were aligned to the hg19 genome of Gencode (release 19) using STAR (v. 2.7.9a) with the previously described parameters. Subsequently, a counts table was generated using the featureCounts tool^78^ and normalization was performed using the DESeq2 R/Bioconductor package^77^. Differential gene expression analysis was then carried out.

### Integrative analysis of ENCODE data

rMATS JunctionCountsOnly files for HepG2 and K562 cells with knockdown of 68 and 70 different splicing factors, respectively, were obtained from the ENCODE portal (https://www.encodeproject.org/). Significantly alternatively spliced events under SF3B1 knockdown were defined with a FDR threshold < 0.05 and |dpsi| > 5%. For this set of splicing events, dpsi values for all other samples with different splicing factor knockdowns were calculated. The dpsi values were then centered and scaled (Z-score) for each splicing event in the formed set. Principal Component Analysis (PCA) was performed using the FactoMineR R/Bioconductor package.

### Cell cultures

Human cancer cell lines SKOV3 (ATCC, HTB-77), MESOV (ATCC, CRL-3272), TOV-112D (ATCC, CRL-3593), TOV-21G (ATCC, CRL-11730), A2780 (Sigma, 93112519), HEY (Cellosaurus, CVCL_0297), Hep G2 (ATCC, HB-8065), HT-29 (ATCC, HTB-38), MDA-MB-231 (ATCC, HTB-26) and mouse ovarian surface epithelial cell line ID8 (Sigma, SCC145) were cultured as adherent monolayers in DMEM medium supplemented with 10% FBS (Invitrogen, #A3382001), 2 mM L-glutamine, and 1% penicillin/streptomycin. Human ovarian cancer cell line OVCAR3 (ATCC, HTB-161) and murine colon carcinoma cell line CT26 (ATCC, CRL-2638) were grown in RPMI medium supplemented with 10% FBS, 2 mM L-glutamine, and 1% penicillin/streptomycin. hTERT-immortalized cells that were isolated from the fallopian tube of a female donor FT282 (ATCC, CRL-3449) was grown as adherent monolayers in DMEM medium supplemented with 10% FBS, 2 mM L-glutamine, and 1% penicillin/streptomycin. Primary culture of human dermal fibroblasts was grown in DMEM medium supplemented with 10% FBS, 1% non-essential amino acids, 2 mM L-glutamine, and 1% penicillin/streptomycin. Ethical approval for the fibroblast cell line was obtained from the Research and Clinical Center of Physical-Chemical Medicine. All cell lines were incubated at 37°C in a humidified atmosphere containing 5% CO2.

### Generation of resistant SKOV3 cells

We generated cisplatin-resistant SKOV3 cells from the parental SKOV3 cell line through a six-month protocol. Initially, cells were cultured at IC10 cisplatin concentration for 48 hours, followed by 72 hours in normal medium to enable the growth of surviving cells. This process was iteratively conducted. We gradually escalated cisplatin concentrations to IC15, IC20, and eventually reaching IC50 after six months.

Similarly, pladienolide-B-resistant SKOV3 cells from the parental SKOV3 cell line were generated over four months. Initially cultured at IC10 Pl-B concentration, cells were continuously cultured at IC15. Pl-B concentrations were subsequently elevated to IC20, culminating in the ability to culture cells at the IC50 concentration for the parental cell line.

### Experimental design

#### For RNA sequencing

Experiment 1: SKOV3 cells were treated with a high (100 nM) or low (1.56 nM) dose of splicing inhibitor (pladienolide B) for 24 or 48 hours, after that total RNA was isolated. As a control, SKOV3 cells were incubated with DMSO, with the same volume as pladienolide B, since Pl-B is dissolved in DMSO.

Experiment 2: Types of treatment: “CP” - SKOV3 cells were treated with cisplatin (10 µM) for 12 hours. “Pl-B + CP” - SKOV3 cells were pretreated with pladienolide B (1.56 nM) for 48 hours followed by cisplatin (10 µM) treatment for 12 hours. “Pl-B release” - SKOV3 cells were pretreated with pladienolide B (1.56 nM) for 48 hours and then cultivated in fresh medium without a drug for 12 hours. “DMSO” - SKOV3 cells were pretreated with DMSO (with the same volume as pladienolide B) for 48 hours and then cultivated in fresh medium without DMSO for 12 hours. After indicated SKOV3 treatment regimens, total RNA from each samples was isolated.

Experiment 3: Types of treatment: “CP” - SKOV3 cells were treated with cisplatin (10 µM) for 12 hours. “H3B-8800+cisplatin” - SKOV3 cells were pretreated with H3B-8800 (50 nM) for 48 hours followed by cisplatin (10 µM) treatment for 12 hours. “H3B-8800 release” - SKOV3 cells were pretreated with H3B-8800 (50 nM) for 48 hours and then cultivated in fresh medium without a drug for 12 hours. “DMSO” - SKOV3 cells were pretreated with DMSO (with the same volume as H3B-8800) for 48 hours and then cultivated in fresh medium without DMSO for 12 hours. After indicated SKOV3 treatment regimens, total RNA from each samples was isolated.

#### For proteomic analyses

Experiment 1: SKOV3 cells were treated with a low (1.56 nM) dose of splicing inhibitor pladienolide B for 24 or 48 hours. As a control, SKOV3 cells were incubated with DMSO (with the same volume as pladienolide B), since Pl-B is dissolved in DMSO. After indicated regimens а SKOV3 treatment, cells were lysed and subjected to proteomic analysis.

Experiment 2: Types of treatment: “CP” - SKOV3 cells were treated with cisplatin (10 µM) for 24 hours. “Pl-B + CP” - SKOV3 cells were pretreated with pladienolide B (1.56 nM) for 48 hours followed by cisplatin (10 µM) treatment for 24 hours. “DMSO” - SKOV3 cells were pretreated with DMSO (with the same volume as pladienolide B) for 48 hours and then cultivated in fresh medium without DMSO for 24 hours. After the indicated regimens of SKOV3 treatment, cells were lysed and subjected to proteomic analysis.

Experiment 3: Types of treatment: “CP” - SKOV3 cells were treated with cisplatin for 24 hours. “H3B-8800+cisplatin” - SKOV3 cells were pretreated with H3B-8800 (50 nM) for 48 hours followed by cisplatin (10 µM) treatment for 24 hours. “DMSO” - SKOV3 cells were pretreated with DMSO (with the same volume as H3B-8800) for 48 hours and then cultivated in fresh medium without DMSO for 24 hours. After the indicated regimens of SKOV3 treatment, cells were lysed and subjected to proteomic analysis.

### RNA Isolation

For RNA sequencing, total RNA was isolated from SKOV3 ovarian cancer cells subjected to treatment with a splicing inhibitor, cisplatin, or their combination, as outlined in the “Experimental design” section. The RNeasy Mini Kit (Qiagen, #74104) was employed for total RNA isolation. Subsequently, all samples underwent DNase treatment using the TURBO DNA-free kit (Thermo Scientific, #AM2238) in 50 µl volumes. RNA cleanup was executed using the Agencourt RNA Clean XP kit (Beckman Coulter, A66514). The concentration and quality of the total RNA were assessed using the Quant-it RiboGreen RNA assay (Thermo Scientific, #R11490) and the RNA 6000 Pico kit (Agilent Technologies, #5067-1513), respectively.

### RNA Sequencing

Polyadenylated RNA enrichment and library preparation were conducted using the NEBNext Poly(A) mRNA Magnetic Isolation Module and NEBNext Ultra II Directional RNA Library Prep Kit (NEB, #E7490S and #E7760S), respectively, following the manufacturer’s protocol. A final cleanup of the library was performed using the Agencourt AMPure XP system (Beckman Coulter, #A63882). The size distribution and quality of the libraries were evaluated using a high sensitivity DNA kit (Agilent Technologies #5067-4626). Quantification of libraries was carried out with the Quant-iT DNA Assay Kit, High Sensitivity (Thermo Scientific, #Q33120). Finally, equal quantities of all libraries (12 pM) were subjected to high-throughput sequencing on the Illumina HiSeq 2500 platform, employing 2 × 100 bp paired-end reads (a sequencing depth of approximately 20 million reads per sample) and a 5% Phix spike-in control.

The raw sequence data and processed data have been deposited in the Gene Expression Omnibus under accession number: GSE227111.

### Protein Concentration Determination

Protein concentrations were determined using Quick Start Bradford Protein Assay (Bio-Rad, #5000201) according to the manufacturer’s standard protocol. Bovine Serum Albumin (BSA) served as the standard for calibration.

### In-solution Trypsin Digestion

SKOV3 cells, treated with the splicing inhibitor, cisplatin, or their combination (as detailed in the “Experimental design” section), were harvested, washed three times with PBS, and lysed using a solution of 4% SDS and 50 mM TRIS-HCl (pH=8) containing protease inhibitors (Sigma-Aldrich) as described previously^79^. The cell lysates underwent sonication on ice through 3 cycles of 10 s on/off pulses with a 30% amplitude. After precipitation of proteins with methanol/chloroform, the semi-dry protein pellet was reconstituted in 50 µL of solubilization buffer (8 M urea, 2 M thiourea, 10 mM TRIS-HCl, pH=8). Protein concentration was measured, and 100 μg of each sample underwent in-solution protein digestion. Disulfide bonds were reduced with DTT (final concentration 5 mM) for 30 min at room temperature (RT). Following this, iodoacetamide was added to a final concentration of 10 mM, with samples incubated in the dark at RT for 20 min. The reaction was halted by adding DTT to a final concentration of 5 mM. Samples were diluted with ammonium bicarbonate solution to reduce urea concentration to 2 M. Trypsin (Promega, #V511A) was added at a ratio of 1:100 (w/w), and samples were incubated for 14 h at 37°C. The reaction was stopped by adding trifluoroacetic acid (TFA) to a final concentration of 0.5%. Finally, tryptic peptides were desalted using SDB-RPS membrane (Sigma-Aldrich, 66886-U), vacuum-dried, and stored at −80°C before LC-MS/MS analysis. Before LC-MS/MS analysis, samples were reconstituted in a solution of 5% acetonitrile (ACN) with 0.1% TFA and sonicated.

### LC-MS/MS Analysis

Proteomic analysis of SKOV3 cells was performed using Q Exactive HF mass-spectrometer. Samples were loaded onto 50-cm columns packed in-house with C18 3 μM Acclaim PepMap 100 (Thermo Scientific), using Ultimate 3000 Nano LC System (Thermo Scientific), coupled to the Q Exactive HF mass-spectrometer (Thermo Scientific). Peptides were loaded onto the column, thermostatically controlled at 40°C in buffer A (0.2% Formic acid), and eluted with a linear (120 min) gradient of 4–55% buffer B (0.1% formic acid, 80% ACN) in buffer A at a flow rate of 350 nl/min. Mass-spectrometry data were stored during the automatic switching between MS1 scans and up to 15 MS/MS scans (topN method). The target value for MS1 scanning was set to 3×10^6^ in the range 300–1,200 m/z with a maximum ion injection time of 60 ms and a resolution of 60,000. Precursor ions were isolated at a window width of 1.4 m/z and a fixed first mass of 100.0 m/z. Precursor ions were fragmented by high-energy dissociation with a normalized collision energy of 28 eV. MS/MS scans were saved with a resolution of 15,000 at 400 m/z and at a value of 1×10^5^ for target ions in the range of 200–2,000 m/z with a maximum ion injection time of 30 ms.

### Protein Identification and Spectral Counting

Raw LC-MS/MS data from the Q Exactive HF mass spectrometer underwent conversion to mgf peaklists using MSConvert from the ProteoWizard Software Foundation. The specific command line parameters employed for this process were “--mgf --filter peakPicking true [1,2]”.

For comprehensive protein identification, the generated peaklists were processed through MASCOT (version 2.5.1, Matrix Science Ltd.) and X! Tandem (ALANINE, 2017.02.01, The Global Proteome Machine Organization). The searches were conducted against the UniProt Knowledgebase (taxon human) with a concatenated reverse decoy dataset. The precursor and fragment mass tolerance were set at 20 ppm and 50 ppm, respectively, for both search algorithms. The database search parameters included tryptic digestion with one possible missed cleavage, static modification for carbamidomethyl (C), and dynamic/flexible modification for oxidation (M). X! Tandem’s parameters enabled the rapid detection of protein N-terminal acetylation, peptide N-terminal glutamine ammonia loss, or peptide N-terminal glutamic acid water loss. To compare results and compile the final list of identified proteins, files from both MASCOT and X! Tandem were subjected to Scaffold 5 (version 5.1.0, Proteome Software Inc) for validation and further analysis. The local false discovery rate scoring algorithm with standard experiment-wide protein grouping was employed. For the evaluation of peptide and protein hits, a false discovery rate (FDR) less than 5% was selected for both. FDR estimates were based on reverse decoy database analysis.

### Detection of Splice Isoforms via RT-PCR

For the semiquantitative RT-PCR analysis of mRNA splice isoforms, total RNA was isolated from SKOV3 cells treated according to the specified experimental design for RNA sequencing. The RNeasy Mini Kit (Qiagen, #74104) was utilized following the manufacturer’s instructions, including on-column DNase I digestion (Qiagen, #79254). The RNA concentration was determined using the Infinite 200 PRO microplate plate reader (Tecan). Subsequently, cDNA was synthesized using Magnus Reverse transcriptase (Evrogen, #SK006S) according to the manufacturer’s protocol. PCR reactions were conducted in duplicate with the Encyclo Plus PCR kit (Evrogen, # PK101) on a T100 Thermal Cycler (Bio-Rad) using the following cycling parameters: 3 min at 95°C; 35 cycles x (10 s at 95°C, 20 s at 60°C, 30 s at 72°C); and 3 min at 72°C. The PCR products were analyzed on a 1.8% agarose TBE-gel with GeneRuler 100 bp Plus DNA Ladder for sizing (Thermo Scientific, #SM0322). The intensity of the bands reflected the expression level of the splice isoforms in the original RNA. The PCR samples were also analyzed by Agilent 2100 Bioanalyzer system (Agilent Technologies) with Agilent DNA 1000 Kit (Agilent Technologies, #5067-1504). GAPDH was used as the reference gene. The following splicing event primers were used:

CHEK2-exon8-F: ctattggttcagcaagagaggc and CHEK2-exon8-R: ccccaccactttgtcaaacag; CHEK2-exon10-F: ctgtttgacaaagtggtgggg and CHEK2-exon10-R: aactcctaaactccagcagtcca; RAD51-exon4-F: agcttgaagcaaatgcagatactt and RAD51-exon4-R: gtgatagatccagtctcaattccac; RAD51-exon6-F: gtggaattgagactggatctatcac and RAD51-exon6-R: gggtctggtggtctgtgttga; RIF1-exon23-F: caactttaacaagacctctggcat and RIF1-exon23-R: taactcttcaggataaaccaacatca; RIF1-exon32-F: tgttaataaggttcgccgtgtct and RIF1-exon32-R: cactttctgttgggcatggta; XPA-exon3-F: tgtaaaagcagccccaaagat and XPA-exon3-R: cctgtcggacttcctttgct; GAPDH-F: aatcccatcaccatcttcca and GAPDH-R: tggactccacgacgtactca.

### Western Blotting

SKOV3 cells treated according to the “Experimental design” section (for proteomic analyses) were lysed for 15 min at 4°C in RIPA buffer containing 1% protease inhibitor mix (Cytiva, 80-6501-23). Lysates were pre-cleaned by centrifugation at 16,000 g for 15 min at 4°C. Protein concentrations were determined by BCA assay (Thermo Scientific, 23225). Equal amounts of protein samples (15 μg/lane) were subjected to analysis by 10% SDS-PAGE and transferred to a PVDF membrane (Bio-Rad, 1620264) using either semi-dry or wet electroblotting (for proteins with a molecular weight of 100 kDa and above). The membrane was blocked with 5% nonfat dry milk in PBST for 1 h and then incubated overnight with primary antibodies against FANCI (ABclonal, A3447), DDB2 (ABclonal, A11615), RAD51 (ABclonal, A2829), CHK2 (ABclonal, A19543), LIG1 (Abcam, ab177946). Subsequently, the membrane underwent incubation with peroxidase-conjugated secondary antibodies (Invitrogen, G-21234) for 1 h. Antibodies against GAPDH (Cloud-Clone Corp., PAB932Hu01) and TUBb (Cloud-Clone Corp., PAB870Hu01) served as loading controls. Immunolabeled proteins were detected using a chemiluminescence detection system ChemiDoc (Bio-Rad, USA).

### Survival Assay and Cell Cycle Analysis

Cancer cells were seeded at a density of 5,000 cells per well in 96-well plates, allowed to adhere overnight, and subsequently treated with pladienolide B (Santa Cruz Biotechnology, Inc., USA, cat. #sc-391691) or DMSO (Paneko, F135), along with varying concentrations of cisplatin (Teva, 115849), carboplatin (Teva, Israel), doxorubicin (Teva, Israel), etoposide (Sigma, cat. #341205), or gemcitabine (Ebewe Pharma, 70006711) for 48 hours (simultaneous treatment). For sequential treatment, cells were preincubated with pladienolide B or DMSO for 48 hours, followed by treatment with different concentrations of cisplatin, carboplatin, doxorubicin, etoposide, or gemcitabine for an additional 48 hours. Cell viability was assessed using the MTT reagent (Sigma-Aldrich, M5655) and measured with the AMR-100 Microplate Reader (Allsheng, China). All experiments were performed at least 3 times.

The synergy of drug combinations was evaluated using the Zero-Interaction Potency (ZIP) method with the SynergyFinder R/Bioconductor package. The input cell viability data was reshaped and pre-processed using the ReshapeData function with recommended parameters: impute = TRUE, impute_method = NULL, noise = TRUE. Combinations of drugs with a ZIP synergy score exceeding 10 were considered synergistic.

Caspase 3/7 activity was assessed using the CellEvent Caspase-3/7 Green Flow Cytometry Assay Kit (Thermo Scientific, #C10427) according to the manufacturer’s protocol. Analysis was performed on a NovoCyte Flow Cytometer (ACEA Biosciences, San Diego, USA).

For cell cycle analysis, cells were fixed in 300 μl ice-cold 70% EtOH, while vortexing slightly, incubated at −20°C for at least 2 hours, washed with 3 ml of PBS, and incubated with 0.1% Triton X-100 in PBS supplemented with 4′,6-diamidino-2-phenylindole (DAPI) (1 μg/ml) (Invitrogen, # 62248) for 30 min. Flow cytometry was conducted on a NovoCyte Flow Cytometer with NovoExpress software (ACEA Biosciences, San Diego, USA).

### Single-Cell Gel Electrophoresis (Comet Assay)

SKOV3 cells were treated with pladienolide B, cisplatin, or their combination for 24 hours. Subsequently, 3,000 cells underwent single-cell gel electrophoresis, adhering to Trevigen’s instructions (catalog # 4250-050-K). Images of the resulting comets were captured using a Nikon Eclipse Ts2 microscope (Nikon, Japan), and subsequent quantification was conducted using the CometScore 2.0 software.

### Transient Transfection

To evaluate NMD (nonsense-mediated mRNA decay) activity, SKOV3 cells were seeded onto 6-well plates and transfected with plasmids pNMD+ or pNMD-31. The transfection was carried out using Lipofectamine 3000 transfection reagent (Invitrogen, cat.# L3000015) in accordance with the manufacturer’s protocol. After 48 hours of transfection, the cells underwent treatment with either 1.56 or 3 nM of pladienolide B for 24 hours. Subsequently, the cells were subjected to flow cytometry analysis. The stained samples were then analyzed using the NovoCyte Flow Cytometer (ACEA Biosciences), and the data were processed using NovoExpress.

### Flow Cytometry

After washing the cells with PBS, they were fixed using 4% paraformaldehyde (PFA) solution in PBS. Subsequently, the cells were blocked with serum-free protein block solution (Dako, X0909). A permeabilization step was performed using 0.1% Triton X-100 in water for 30 minutes. Following this, the cells were incubated with primary antibodies overnight. The primary antibodies used were anti-ATM (Invitrogen #MA1-23152), anti-ATM (phospho S1981) (Abcam, ab81292), anti-ATR (Invitrogen #MA1-23158), or phospho-ATR (Thr1989) (Invitrogen, #PA5-77873). After the primary antibody incubation, a 2-hour incubation with Alexa Fluor 488-conjugated secondary antibodies (Thermo Scientific, #A-11008, #A-11001) followed. The stained samples were then analyzed using the NovoCyte Flow Cytometer (ACEA Biosciences), and the data were processed using NovoExpress.

### Fluorescence Microscopy

To prepare for immunofluorescence, SKOV3 cells on chamber slides were fixed using CSK buffer^80^ for 15 minutes. Subsequently, the cells were washed thrice with PBS and then incubated with primary antibodies overnight. The primary antibodies used were anti-gamma H2A.X (phospho S139) (Sigma-Aldrich, #05-636), anti-RPA32/RPA2 (phospho S33) (Abcam, ab21187), DNA-RNA hybrid (Sigma-Aldrich, MABE1095), and anti-DNA PKcs (phospho S2056) antibody (Abcam, ab18192). After the primary antibody incubation, a 2-hour incubation with Alexa Fluor 488-conjugated secondary antibodies (Thermo Scientific, #A-11008, #A-11001) followed. Subsequently, the cells were incubated with DAPI solution for 20 min. The slides were mounted with Fluoroshield mounting medium (Sigma-Aldrich) and covered with cover glasses. Images were captured using the Nikon Eclipse Ni-E microscope in DAPI, FITC, and TRITC channels. Image analysis was performed using ImageJ software with FindFoci plugins.

### Animal Studies in vivo and ex vivo

All experimental procedures involving animals were conducted in compliance with ethical standards and were approved by the Institutional Animal Care and Use Committee (IACUC) of the Shemyakin-Ovchinnikov Institute of Bioorganic Chemistry Russian Academy of Sciences, in accordance with the IACUC protocol #299 (1 January 2020 – 31 December 2022) and in accordance with the Declaration of Helsinki and approved by Local Ethics Committee of Federal State Budgetary Educational Institution of Higher Education “Privolzhsky Research Medical University” of the Ministry of Health of the Russian Federation (protocol # 10, 26.06.2020). For the *in vivo* experiments, CT26 cells stably expressing near-infrared fluorescent protein eqFP650 (FP731, Evrogen) were injected subcutaneously into female Balb/c mice (12-14 weeks old) in the amount of 250,000 cells in PBS. Tumor growth was monitored thrice a week for 3.5 weeks. Tumor volumes were calculated using the formula V = (length × width^2) × 3.14/6. On the day 10 after cancer cells injection, when the tumor diameter reached ∼5-7 mm, therapy was started. Mice were randomized into treatment cohorts, and drugs were administered intraperitoneally with the following group assignments (n = 5-10 per group): “Pl-B + CP”: Mice treated with a combination of pladienolide B (1 mg/kg) and cisplatin (5 mg/kg or 2.5 mg/kg). Cycles consisted of injections of pladienolide B followed by cisplatin after 24 hours, repeated four times with 1 or 2 days of rest between cycles during 2 weeks (totaling 4 injections of Pl-B and 4 injections of cisplatin); “CP”: Mice treated with cisplatin (5 mg/kg or 2.5 mg/kg) twice a week during 2 weeks (totaling 4 injections of cisplatin); “Pl-B”: Mice treated with pladienolide B (1 mg/kg) twice a week during 2 weeks (totaling 4 injections of Pl-B); “Control”: Untreated mice injected with saline solution instead of drugs twice a week during 2 weeks. On day 25 of tumor growth whole-body fluorescence visualization of mice was performed on a LumoTrace FLUO bioimaging system (Abisense, Russia) and IVIS Spectrum (Caliper Life Sciences, USA). Shaved animals were anesthetized and imaged using an excitation 590-nm diode and 655-nm long-pass filter with exposure time 3,000 ms for LumoTrace FLUO and using an excitation 605-nm and 660-nm long-pass filter with exposure time 30 ms for IVIS Spectrum.

Following euthanasia on day 25, each tumor was dissected, fixed in 10% neutral buffered formalin for 24 hours, embedded in paraffin, and sectioned for histopathologic examination. Four-micrometer sections of tissues were stained with hematoxylin and eosin. Tumor structure images were captured using the Leica Aperio AT2 microscope (Leica, Japan) and analyzed with Aperio ImageScope. For counting the number of mitosis, each slice was randomly selected and 12 visual fields per slice were examined. All mitosis figures were counted in each field under a light microscope at x 400 magnification. The mitotic count was expressed as the number of mitoses per mm2 of the tumor.

### Quantification and statistical analysis

All data are presented as mean ± SD. The number of replicates for each experiment is specified in the figure legends, indicating independent biological replicates. Statistical analyses were conducted using R or GraphPad PRISM8. Details regarding the specific statistical tests employed and sample sizes for each experiment can be found in the figure legends or the “Methods” section. Sample sizes were determined based on our prior studies and the authors’ expertise^81^. No samples, mice, or data points were excluded from the reported analyses. Unless otherwise mentioned, all western blots, flow cytometry data, and microscopy images represent three biologically independent experiments.

---

**Extended Data Figure 1.**
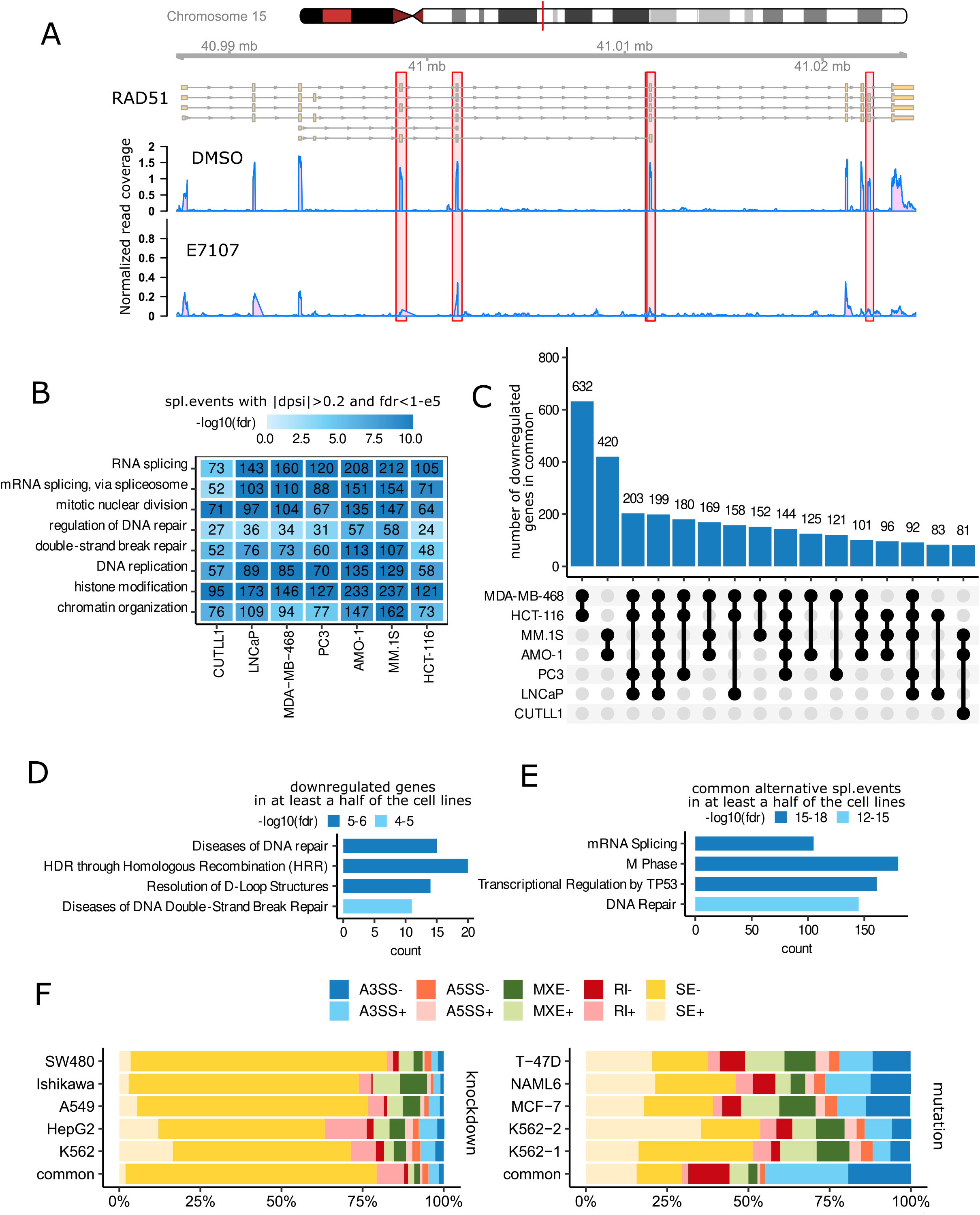
Consistent pre-mRNA splicing and gene expression patterns Induced by a common splicing Inhibitor across different cancer cell lines. **A** – RNA-seq read coverage for the RAD51 gene. Y-axis represents read-depth normalized density from one MM.1S cell line replicate after E7107 treatment. Coverage alterations are observed in exons, not introns, under splicing inhibitors. **B** – Heat map of Reactome pathway enrichment analysis, highlighting common pathways in 7 cancer cell lines for genes undergoing splicing changes after E7107 treatment. Selected splicing events: FDR < 1-e5, |dpsi| > 0.2. X-axis: Cancer cell line names; Y-axis: Pathway names. Numbers in boxes represent number of genes associated with the pathway for the cell line, FDR < 0.05. Blue color scale indicates p-value. **С** – UpSet plot indicating the number of common downregulated genes among 7 cancer cell lines after E7107 treatment. **D** – Functional analysis of genes downregulated in over half of the cell lines after E7107 treatment. Details on cell lines and E7107 doses in Table 1. **E** – Functional analysis of genes exhibiting common differential splicing events in over half of the cell lines after E7107 treatment. Details on cell lines and E7107 doses in Table 1. **F** – Stacked bar plot displaying the percentage of each alternative splicing event type among all differential splicing events in cancer cell lines with SF3B1 mutation or after SF3B1 knockdown. Splicing events: SE - skipped exon, A5SS - alternative 5′ splice site, A3SS - alternative 3′ splice site, RI - retained intron, MXE - mutually exclusive exons. Plus sign: dpsi > 0.05, Minus sign: dpsi < −0.05.

**Extended Data Figure 2.**
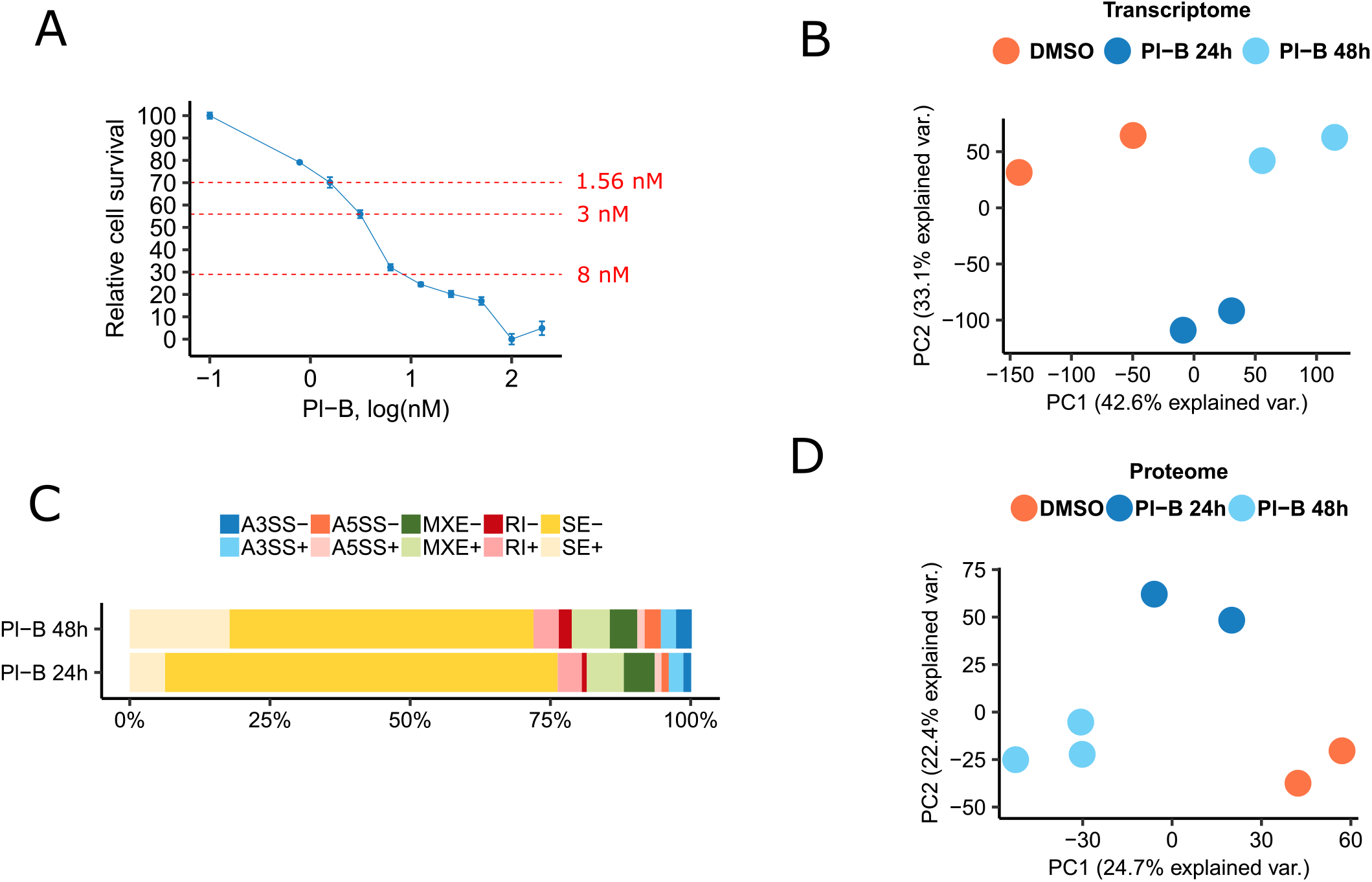
A rise in the expression of splicing factors leads to exon skipping after pladienolide B treatment. **A** – Dose-response curves obtained by MTT assay of SKOV3 cells treated with different concentrations of pladienolide B for 48 hours. Each data point represents mean values ± SD (n = 3). IC30, IC45 and IC70 values are determined by fitting a normalized model to data with nonlinear regression using GraphPad Prism software. **B** – Principal component analysis (PCA) of log2-transformed normalized gene expression values comparing control (DMSO) and treated samples. Each point represents the expression profile of one sample. Types of treatment: “DMSO” - cells treated with DMSO for 24 hours, “Pl-B 48h” - cells treated with pladienolide B for 48 hours (1.56 nM), “Pl-B 24h” - cells treated with pladienolide B for 24 hours (1.56 nM). **C** – Stacked bar plot displaying the percentage of each alternative splicing event type among all differential splicing events in SKOV3 cells after 24 or 48 hours of pladienolide B treatment. Splicing events: SE - skipped exon, A5SS - alternative 5′ splice site, A3SS - alternative 3′ splice site, RI - retained intron, MXE - mutually exclusive exons. Plus sign: dpsi > 0.05, Minus sign: dpsi < −0.05. **D** – Principal component analysis (PCA) of proteomic profiles comparing control (DMSO) and treated samples. Each point represents the proteomic profile of one sample. Types of treatment: “DMSO” - cells treated with DMSO for 24 hours, “Pl-B 48h” - cells treated with pladienolide B for 48 hours (1.56 nM), “Pl-B 24h” - cells treated with pladienolide B for 24 hours (1.56 nM).

**Extended Data Figure 3.**
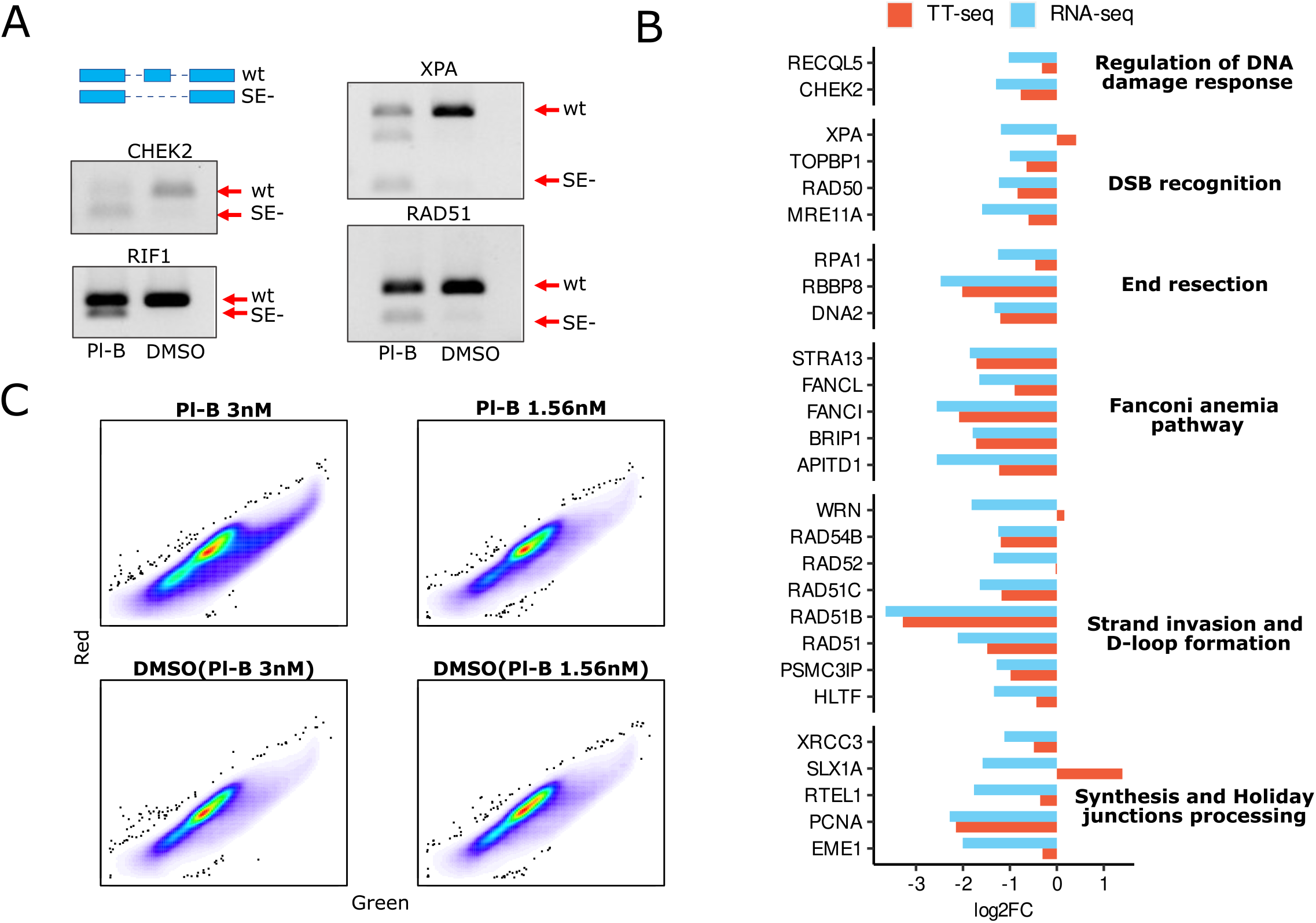
Impact of SF3B1 inhibition on NMD activity. **A** – RT-PCR results confirming exon skipping events in *XPA*, *RAD51*, *RIF1*, and *CHEK2* mRNAs in SKOV3 cells treated with 1.56 nM pladienolide B for 24 hours. “wt” indicates the wild-type transcript, “SE-” indicates the transcript with an exon skipping event. **B** – Bar graph comparing log2-fold expression changes of selected DNA repair genes identified by RNA-seq and TT-seq data analysis in the CUTLL1 cell line after 24 hours of E7107 exposure. **C** – Flow cytometry analysis dot plots (in green and red channels) of SKOV3 cells transiently transfected with two fluorescent proteins carried on a plasmid: one (green) is encoded by the NMD-targeted transcript, and the other (red) serves as an expression efficiency control. Flow cytometry analysis performed for SKOV3 cells treated with different doses of pladienolide B or DMSO for 24 hours.

**Extended Data Figure 4.**
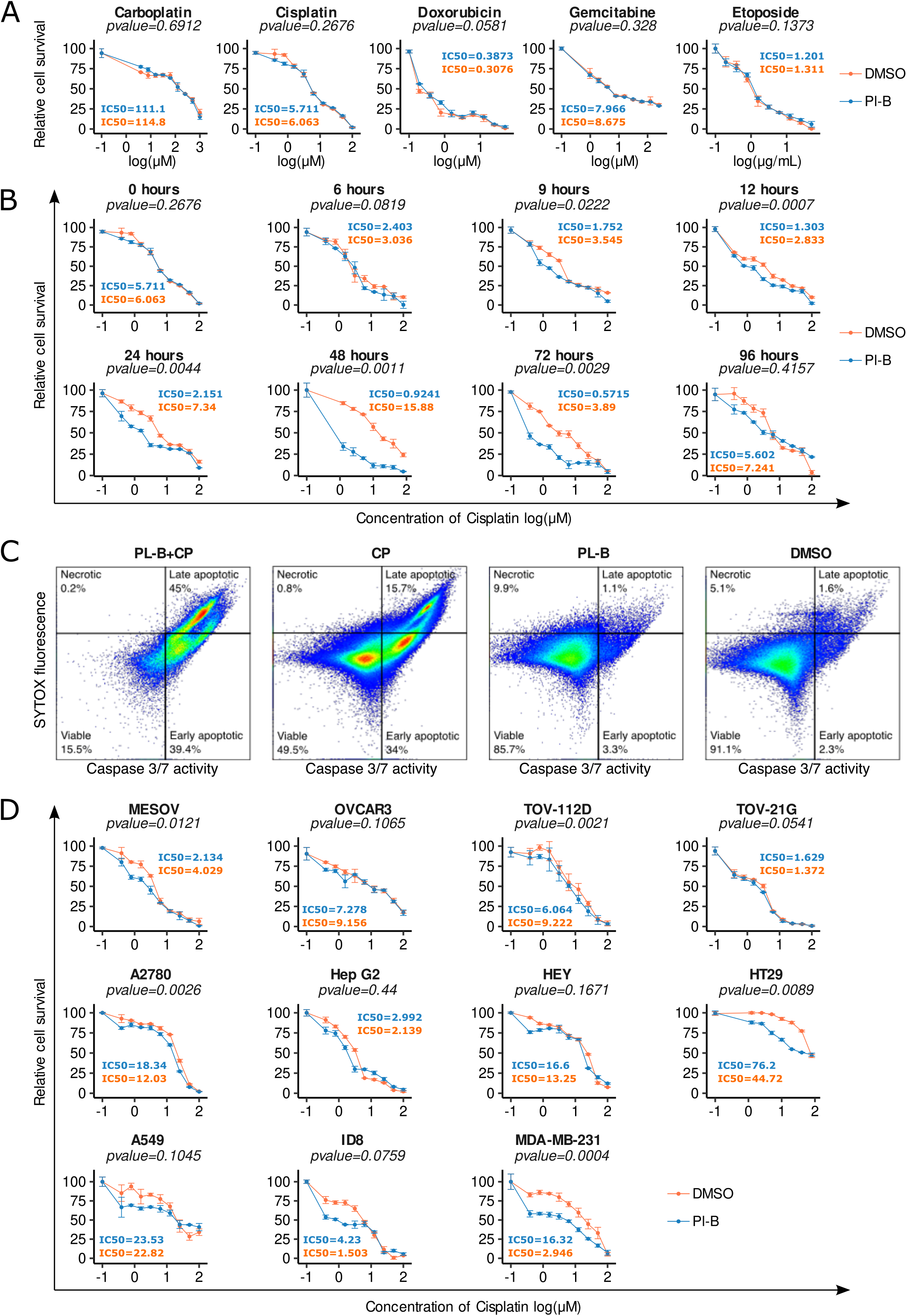
In vitro impact of simultaneous and sequential combination therapy with pladienolide B and cisplatin. **A** – Dose-response curves from MTT assays of SKOV3 cells treated with a fixed concentration of pladienolide B (1.56 nM) and various concentrations of carboplatin, cisplatin, doxorubicin, etoposide, or gemcitabine simultaneously for 48 hours. DMSO was used as a control instead of pladienolide B. **B** – Dose-response curves from MTT assays of SKOV3 cells pretreated with 1.56 nM pladienolide B or DMSO (control) for different durations (0, 6, 9, 12, 24, 48, 72, or 96 hours) followed by treatment with different concentrations of cisplatin for 48 hours. **C** – CellEvent Caspase-3/7 Green Flow Cytometry Assay of SKOV3 cells after different types of treatment. “Pl-B + CP” - pretreatment with pladienolide B (1.56 nM, 48 hours) followed by treatment with cisplatin (10 µM, 24 hours); “CP” - pretreatment with DMSO (48 hours) followed by treatment with cisplatin (10 µM, 24 hours); “Pl-B” - pretreatment with pladienolide B (1.56 nM, 48 hours) followed by fresh medium without pladienolide B (24 hours); “DMSO” - pretreatment with DMSO (48 hours) followed by fresh medium without DMSO (24 hours). **D** – Dose-response curves from MTT assays of various cell lines (MESOV, OVCAR3, TOV112D, TOV21G, A2789, Hep G2, HEY, HT29, A549, ID8, MDA-MB-231) treated with a fixed concentration of pladienolide B (1.56 nM) and different concentrations of cisplatin simultaneously for 48 hours. DMSO served as the control instead of pladienolide B. Each data point in A, B, D represents mean values ± SD (n = 3). IC50 values were determined by fitting a normalized model to data with nonlinear regression using GraphPad Prism software. Significance assessed via a paired, two-tailed Student’s t-test.

**Extended Data Figure 5.**
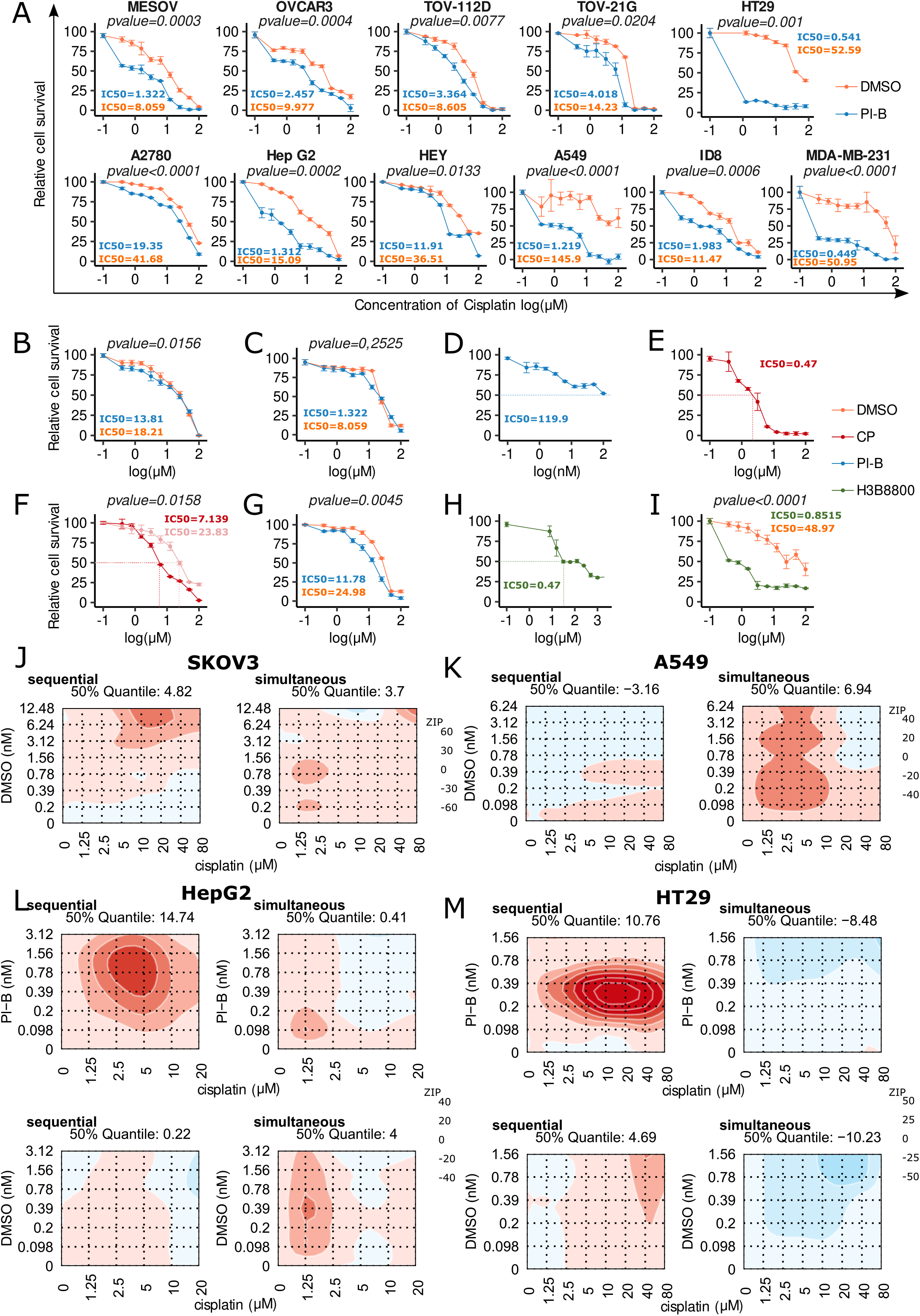
Synergy assessment of pladienolide B and cisplatin combination. **A** – Dose-response curves from MTT assays of various cell lines (MESOV, OVCAR3, TOV112D, TOV21G, A2780, Hep G2, HEY, HT29, A549, ID8, MDA-MB-231) pretreated with 1.56 nM pladienolide B (48 hours) followed by treatment with different concentrations of cisplatin for 48 hours. DMSO was used as the control instead of pladienolide B. **B** – Dose-response curves from MTT assays of human primary fibroblasts pretreated with 1.56 nM pladienolide B (48 hours) followed by treatment with different concentrations of cisplatin for 48 hours. **C** – Dose-response curves from MTT assays of FT282 cells pretreated with 1.56 nM pladienolide B (48 hours) followed by treatment with different concentrations of cisplatin for 48 hours. **D-E** – Dose-response curves from MTT assays of pladienolide B-resistant SKOV3 cells treated with different concentrations of pladienolide B (D) or cisplatin (E) for 48 hours. The method for generating resistant SKOV3 cells is described in the “Methods” section. **F** – Dose-response curves from MTT assays of SKOV3 cells treated with different concentrations of H3B-8800 (another splicing inhibitor) for 48 hours. **G** – Dose-response curves from MTT assays of SKOV3 (red line) and cisplatin-resistant SKOV3 (light red line) cells treated with different concentrations of cisplatin for 48 hours. **H** – Dose-response curves from MTT assays of cisplatin-resistant SKOV3 cells pretreated with 1.56 nM pladienolide B (48 hours) followed by treatment with different concentrations of cisplatin for 48 hours. **I** – Dose-response curves from MTT assays of SKOV3 cells pretreated with 50 nM H3B-8800 (48 hours) followed by treatment with different concentrations of cisplatin for 48 hours. **J-K** – Synergy landscapes for SKOV3 (J) and A549 (K) cells after simultaneous or sequential treatment with different doses of DMSO (y-axis) and cisplatin (x-axis). Zero-Interaction Potency (ZIP) synergy scores were calculated for the combination of DMSO in different volume range (according to Pl-B volume for 0 nM–6.24 nM concentration range) and cisplatin (0 µM–80 µM) for SKOV3 and A549 cells. For simultaneous regimen, сells were treated with DMSO and cisplatin simultaneously (48 hours). For sequential regimen, cells were pretreated with different volumes of DMSO (48 hours) followed by treatment with different concentrations of cisplatin for 48 hours. Cell survival was calculated in comparison to cisplatin untreated cells (DMSO only). ZIP values below 0 indicate antagonism (blue), 0 - 10 indicate additivity (from white to light red), and above 10 (corresponding to a deviation from the reference model above 10%) indicate synergy (dark red). **L-M** – Synergy landscapes for HepG2 (L) and HT29 (M) cells. Each data point represents mean values ± SD (n = 3). IC50 values were determined by fitting a normalized model to data with nonlinear regression using GraphPad Prism software.

**Extended Data Figure 6.**
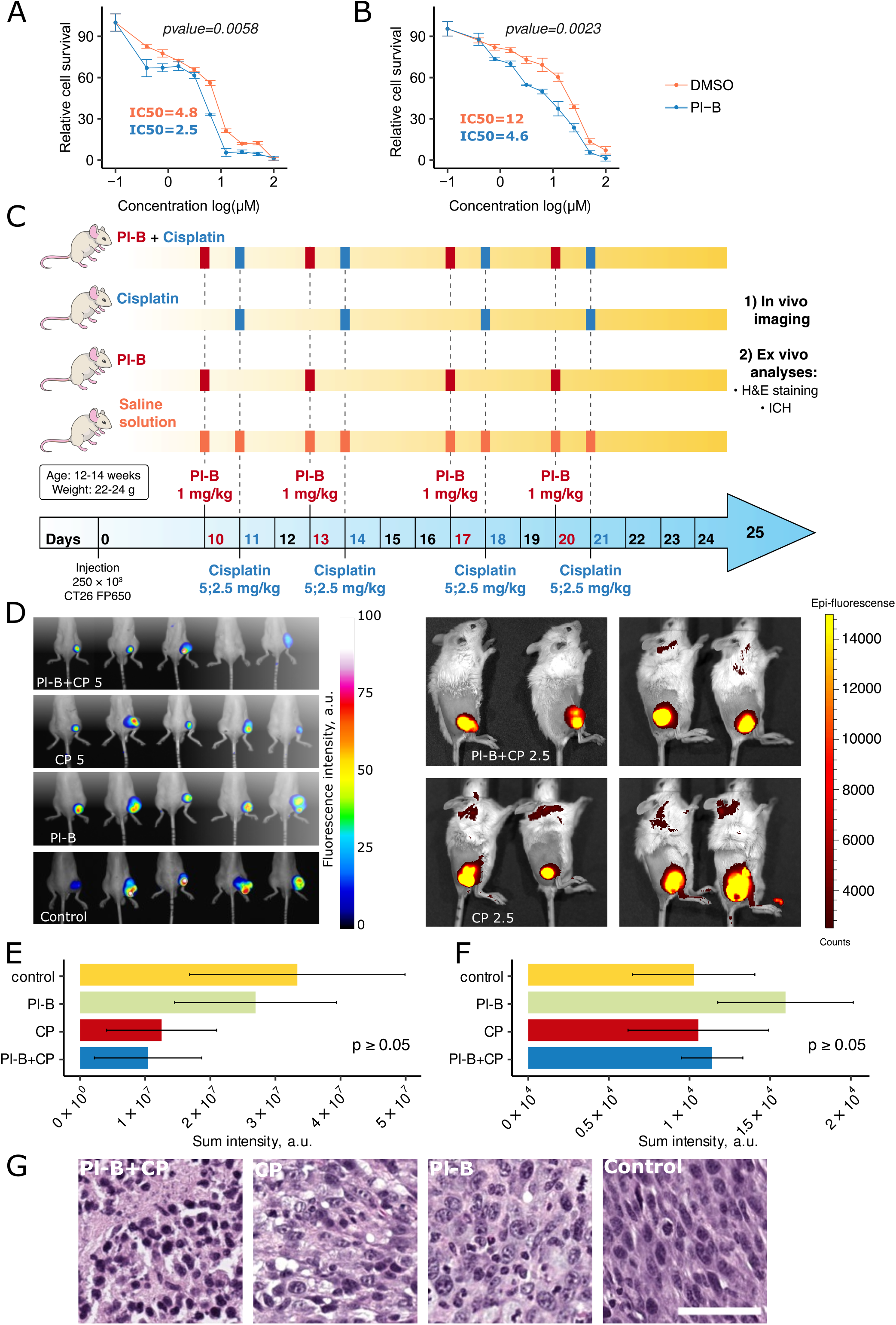
**The effect of combination treatment with pladienolide B and cisplatin on CT26 cells *in vitro* and *in vivo*** **A** – Dose-response curves from MTT assays of CT26 cells with a fixed concentration of pladienolide B (1.56 nM) and varying concentrations of cisplatin simultaneously added for 48 hours. DMSO was used as the control instead of pladienolide B (according to 1.56 nM Pl-B volume). **B** – Dose-response curves from MTT assays of CT26 cells pretreated with 1.56 nM of pladienolide B or DMSO (according to 1.56 nM Pl-B volume) for 48 hours followed by treatment with different concentrations of cisplatin for 48 hours. Each data point on (A) and (B) represents mean values ± SD (n = 3). IC50 values were determined by fitting a normalized model to data with nonlinear regression using GraphPad Prism software. Significance was determined using a paired, two-tailed Student’s t-test. **C** – Scheme of *in vivo* experiment. BALB/c mice were subcutaneously inoculated with eqFP650- expressing CT26 colon carcinoma cells on day 0; on day 10 they were treated either by monotherapy with pladienolide B (1 mg/kg) or cisplatin (5 mg/kg or 2.5 mg/kg), or by combination therapy with the same drugs. Drugs were injected intraperitoneally, days of injection are indicated in figure. **D** – Fluorescence *in vivo* whole-body imaging of eqFP650-expressing tumors in mice on day 25 of tumor growth. The scale bar indicates fluorescence intensity. **E-F** – Columnar graphs representing the sum intensity of eqFP650 fluorescence in CT26 tumors for each treatment group, calculated using Abisense software (E) and Living Image Software (F). Significance was determined using a paired, two-tailed Student’s t-test, mean values ± SD. **G** – Representative microphotographs of hematoxylin and eosin (H&E) staining in CT26 tumors for each *ex vivo* treatment group (original magnification, ×100). According to histopathological analysis, isolated tumors showed a typical hypercellular solid carcinoma phenotype with high grade atypia in the tumor cells and extensive necrotic areas. Scale bar = 50 mm.

**Extended Data Figure 7.**
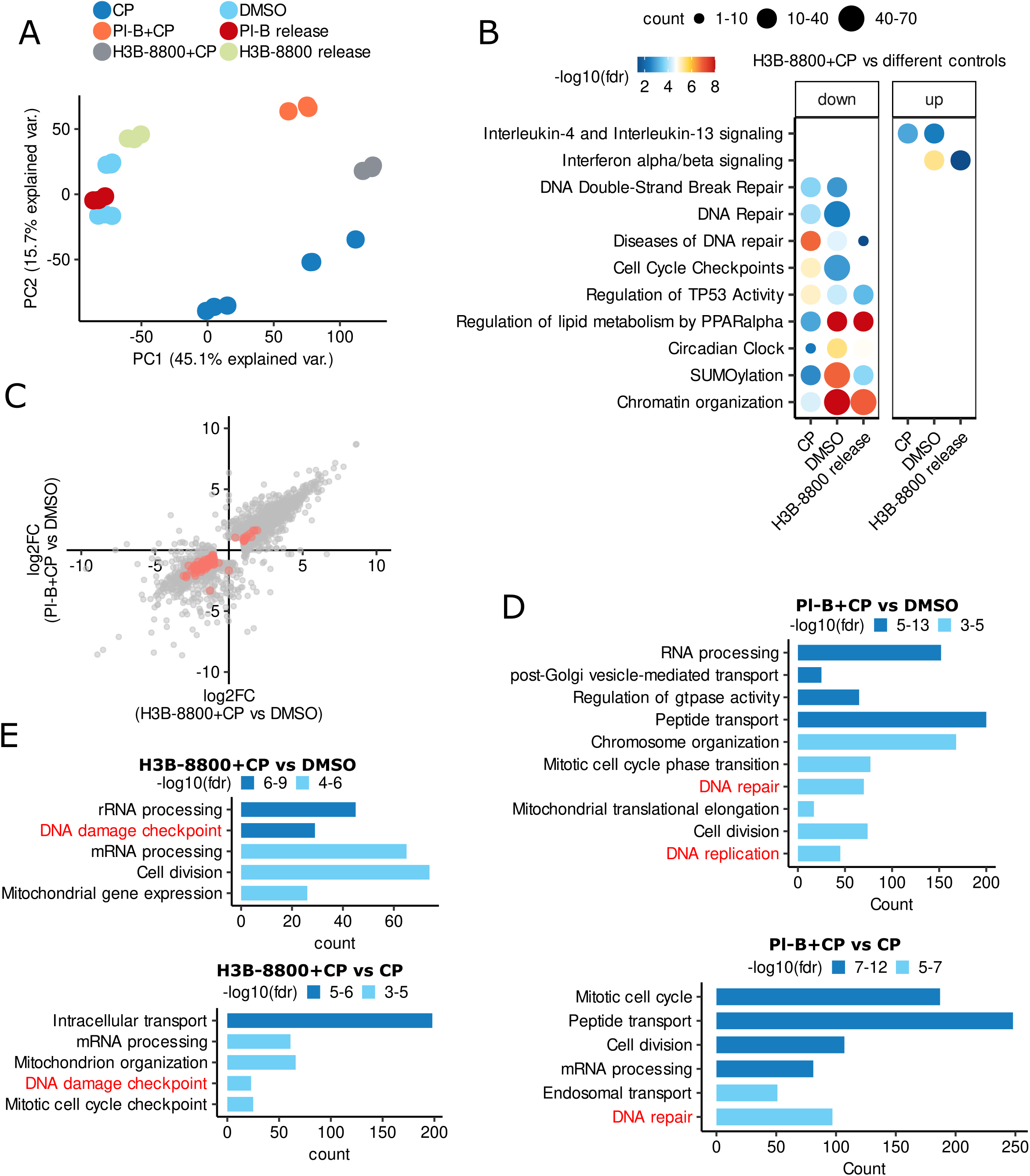
Pretreatment with H3B-8800 also suppresses DNA repair genes. **A** – Principal component analysis (PCA) of the log2-transformed normalized gene expression values between control (DMSO) and treated samples after batch effect removal by ComBat function. Each point represents the expression profile of one sample. Types of treatment: “DMSO” - cells treated with DMSO for 24 hours, “CP” - cells treated with cisplatin for 12 hours, “Pl-B + CP” and “Н3B-8800 + CP” - cells pretreated with pladienolide B (1.56 nM) or H3B-8800 (50 nM) for 48 hours followed by cisplatin (10 µM) treatment for 12 hours, “Pl-B release” and “Н3B-8800 release” - cells treated with pladienolide B (1.56 nM) or H3B-8800 (50 nM) for 48 hours and then cultivated in fresh medium without a drug for 12 hours. **B** – Dot plot showing Reactome pathways (FDR < 0.05) enrichment analysis of differentially expressed genes/proteins in SKOV3 cells treated with the proposed drug combination compared to different controls as described in Extended Data Fig. 7A. The size of the dot is based on gene/protein count enriched in the pathway, and the color of the dot shows the pathway enrichment significance. **C** – Scatter plot of log2-fold changes in gene expression profiles after two types of combination treatment. y-axis represents log2-fold change of gene expression after sequential combination treatment with pladienolide B and cisplatin; x-axis represents the log2-fold change of gene expression after sequential combination treatment with Н3B-8800 and cisplatin. Orange color indicates differentially expressed DNA repair genes. Only genes differentially expressed in at least one of the two comparisons were used for the illustration (FDR < 0.1, |log2-fold change| > 1). **D** – topGO enrichment analysis of downregulated proteins in SKOV3 cells after sequential combination treatment with pladienolide B and cisplatin compared with two control groups: cells after cisplatin or DMSO treatment. **E** – topGO enrichment analysis of downregulated proteins in SKOV3 cells after sequential combination treatment with Н3B-8800 and cisplatin compared with two control groups: cells after cisplatin or DMSO treatment.

**Extended Data Figure 8.**
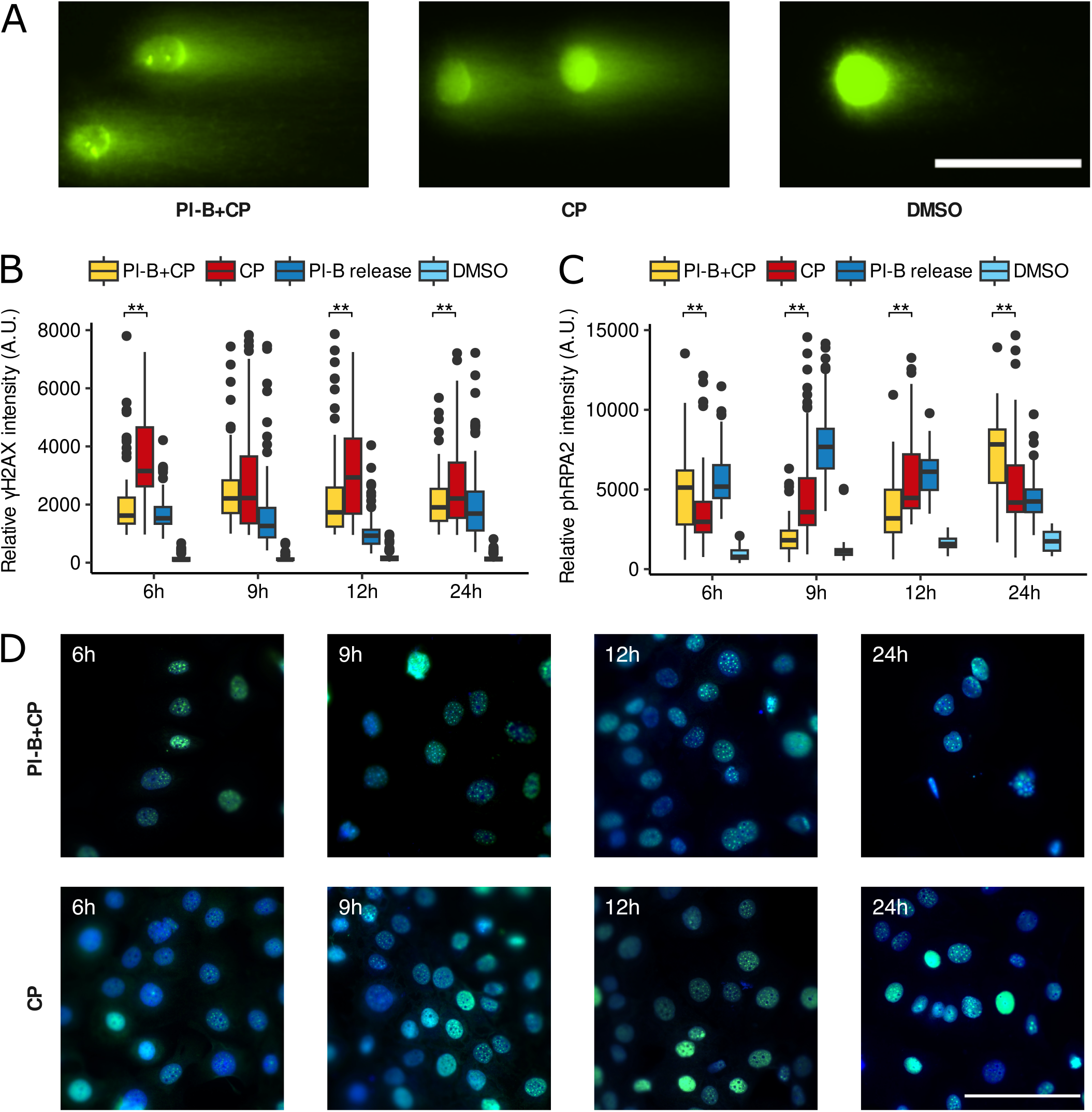
Different effects of combination treatment with pladienolide B and cisplatin on ovarian cancer cells *in vitro*. **A** – Representative comet assay images of SKOV3 cells stained with SYBR green after different treatments and observed under fluorescence microscope. Scale bar = 50 μm. **B** – Box plots showing γH2AX fluorescence intensity in SKOV3 cells after different treatments. The fluorescence intensity was measured using ImageJ software. 80-200 cells were analyzed in each sample. **C** – Box plots showing phRPA2 fluorescence intensity in SKOV3 cells after different treatments. The fluorescence intensity was measured using ImageJ software. 80-200 cells were analyzed in each sample. **D** – Representative immunofluorescence images of SKOV3 cells stained for phosphorylated RPA2 (Ser33, green) and with DAPI (blue) after different treatments. Scale bar = 100 μm. “Pl-B + CP” - SKOV3 cells were pretreated with 1.56 nM of pladienolide B for 48 hours then the medium was changed to the medium with 10 µM of cisplatin for 24 hours (in A) or 6h, 9h, 12h and 24h (in B, C, D); “CP” - SKOV3 cells were incubated with DMSO for 48 hours then the medium was changed to the medium with 10 µM of cisplatin for 24 hours (in A) or 6h, 9h, 12h and 24h (in B, C, D); “Pl-B release” - SKOV3 cells were incubated with 1.56 nM of pladienolide B for 48 hours, then the medium was changed to fresh one without pladienolide B for 6h, 9h, 12h or 24h (in B, C); “DMSO” - SKOV3 cells were incubated with DMSO for 48 hours (in A, D, E) or 6h, 9h, 12h and 24h (in F, G); “Pl-B” - SKOV3 cells were treated with 1.56 nM of pladienolide B for 48 hours (in A) or 6h, 9h, 12h and 24h (in B, C).

## Supplementary Table

**Supplementary Table S1. - An atlas of alternative splicing events caused by perturbations in functional activity of SF3B1**

List1 - A list of common splicing events in 7 cancer lines after E7107 treatment

List2 - The alternative splicing analysis of SKOV3 cells after pladienolide B treatment for 24 or 48 hours

List3 - A list of common splicing events in 4 cancer lines with and without mutation in SF3B1 gene

List4 - A list of common splicing events in 5 cancer lines after SF3B1 knockdown

**Supplementary Table S2 - Transcriptomic profiles of different cancer cell lines after splicing inhibitor exposure, SF3B1 knockdown or with SF3B1 mutation**

List1 - Analysis of differentially expressed genes in 7 cancer cell lines after E7107 exposure List2 - Analysis of differentially expressed genes in SKOV3 cells after pladienolide B treatment for 24 or 48 hours

List3 - Analysis of differentially expressed genes in 5 cancer cell lines after SF3B1 knockdown

**Supplementary Table S3. - Proteomic profiles of SKOV3 cells at different time points after pladienolide B treatment**

List1 - Proteins identified by LC-MS/MS analysis of SKOV3 cells treated or untreated with pladienolide B cells for 24 or 48 hours

**Supplementary Table S4. - A landscape of DNA repair genes dysregulated after splicing inhibitor treatment**

List1 - Analysis of DNA repair genes differentially expressed in 8 different cancer lines treated with splicing inhibitor

**Supplementary Table S5. - Transcriptomic profiles of SKOV3 cells after treatment with splicing inhibitor, cisplatin or their combination**

List1 - Analysis of differentially expressed genes in SKOV3 cells after treatment with pladienolide B, cisplatin or their combination

List2 - Analysis of differentially expressed genes in SKOV3 cells after treatment with Н3B-8800, cisplatin or their combination

**Supplementary Table S6. - Proteomic profiles of SKOV3 cells after treatment with splicing inhibitor, cisplatin or their combination**

List1 - Proteins identified by LC-MS/MS analysis in SKOV3 cells after treatment with pladienolide B, cisplatin or thier combination

List2 - Proteins identified by LC-MS/MS analysis in SKOV3 cells after treatment with Н3B-8800, cisplatin or their combination

